# Principles of RNA processing from analysis of enhanced CLIP maps for 150 RNA binding proteins

**DOI:** 10.1101/807008

**Authors:** Eric L Van Nostrand, Gabriel A Pratt, Brian A Yee, Emily Wheeler, Steven M Blue, Jasmine Mueller, Samuel S Park, Keri E Garcia, Chelsea Gelboin-Burkhart, Thai B Nguyen, Ines Rabano, Rebecca Stanton, Balaji Sundararaman, Ruth Wang, Xiang-Dong Fu, Brenton R Graveley, Gene W Yeo

**Affiliations:** Department of Cellular and Molecular Medicine, University of California San Diego, La Jolla, CA; Institute for Genomic Medicine, University of California San Diego, La Jolla, CA; Department of Genetics and Genome Sciences, Institute for Systems Genomics, UConn Health, Farmington, CT

**Keywords:** eCLIP, CLIP-seq, RNA binding protein, RNA processing

## Abstract

A critical step in uncovering rules of RNA processing is to study the *in vivo* regulatory networks of RNA binding proteins (RBPs). Crosslinking and immunoprecipitation (CLIP) methods enabled mapping RBP targets transcriptome-wide, but methodological differences present challenges to large-scale integrated analysis across datasets. The development of enhanced CLIP (eCLIP) enabled the large-scale mapping of targets for 150 RBPs in K562 and HepG2, creating a unique resource of RBP interactomes profiled with a standardized methodology in the same cell types. Here we describe our analysis of 223 enhanced (eCLIP) datasets characterizing 150 RBPs in K562 and HepG2 cell lines, revealing a range of binding modalities, including highly resolved positioning around splicing signals and mRNA untranslated regions that associate with distinct RBP functions. Quantification of enrichment for repetitive and abundant multi-copy elements reveals 70% of RBPs have enrichment for non-mRNA element classes, enables identification of novel ribosomal RNA processing factors and sites and suggests that association with retrotransposable elements reflects multiple RBP mechanisms of action. Analysis of spliceosomal RBPs indicates that eCLIP resolves AQR association after intronic lariat formation (enabling identification of branch points with single-nucleotide resolution) and provides genome-wide validation for a branch point-based scanning model for 3’ splice site recognition. Further, we show that eCLIP peak co-occurrences across RBPs enables the discovery of novel co-interacting RBPs. Finally, we present a protocol for visualization of RBP:RNA complexes in the eCLIP workflow using biotin and standard chemiluminescent visualization reagents, enabling simplified confirmation of ribonucleoprotein enrichment without radioactivity. This work illustrates the value of integrated analysis across eCLIP profiling of RBPs with widely distinct functions to reveal novel RNA biology. Further, our quantification of both mRNA and other element association will enable further research to identify novel roles of RBPs in regulating RNA processing.

## Background

RNA can act as a carrier of information from the nucleus to the cytoplasm in the processing of protein-coding genes, as a regulatory molecule that can control gene expression, and even as an extracellular signal to coordinate trans-generational inheritance [1–3]. RNA binding proteins (RBPs) interact with RNA through a wide variety of primary sequence motifs and RNA structural elements to control all processing steps [3]. Furthermore, with the increase in the number of RBPs that are becoming associated with human diseases, identifying their RNA targets and how they are regulated has become an unmet, urgent need.

To identify direct RNA targets of RBPs, RNA immunoprecipitation (RIP) and crosslinking and immunoprecipitation (CLIP) methods are frequently used. CLIP-based methods utilize UV crosslinking to covalently link an RBP with its bound RNA in live cells, enabling both stringent immunoprecipitation washes and denaturing SDS-PAGE protein gel electrophoresis and nitrocellulose membrane transfer which serves to remove background unbound RNA [4]. Analyses of single RBP binding profiles by CLIP have provided unique insights into basic mechanisms of RNA processing, as well as identified downstream effectors that drive human diseases [5–7]. Further efforts to profile multiple human RBPs in the same family or regulatory function by CLIP illustrated coordinated and complex auto- and cross-regulatory interactions among RBPs and their targets [8–10]. Rising interest in organizing public deeply sequenced CLIP datasets to enable the community to extract novel RNA biology is apparent from newly available computational databases and integrative methods [11, 12]. However, methodological differences between CLIP approaches, combined with simple experimental variability between labs and variation in acceptable quality control metrics, add significant challenges to interpretation of differences observed.

The field of transcription regulation observed similar challenges and opportunities in integrating transcription factor target profiles [13]. To address this challenge, the ENCODE consortium piloted large-scale profiling of transcription factor targets using a single standardized chromatin immunoprecipitation (ChIP-seq) protocol [14]. The initial effort to profile 119 factors generated a unified dataset for creating and assaying robust quality assessment standards [15], and led to insights into modeling transcription factor complexes, binding modalities, and regulatory networks [16]. More critically, however, this has served as an invaluable resource for researchers to annotate potential functional variants [17] and generate hypotheses across a variety of fields of interest. This success suggested that a similar effort to profile RBP targets using a standardized methodology could similarly drive significant insights in RNA biology.

To this end we introduced the enhanced CLIP (eCLIP) methodology featuring a size-matched input control [18] and characterized hundreds of immunoprecipitation-grade antibodies with a standardized work-flow [19] to generate 223 eCLIP datasets profiling targets for 150 RBPs in K562 and HepG2 cell lines [20]. Along with orthogonal data types, we provided insights into localized RNA processing, studied the interplay between *in vitro* binding motifs and RBP association (and factor-responsive targets) in live cells, and identified novel effectors of RNA stability and alternative splicing [20].

Here, we extend our previous study by providing further insight into how integrative analysis of RBP target profiles by eCLIP can reveal both general principles of RNA processing as well as specific mechanistic insights for individual RBPs. Although most CLIP analysis typically focuses on binding to mRNAs (both intronic and exonic), we find that for 70% of RBPs the dominant enrichment signature is instead a variety of multicopy and non-coding elements (including structural RNAs such as ribosomal RNAs and spliceosomal snRNAs, retrotransposable and other repeat elements, and mitochondrial RNAs). These analyses can be then used to generate hypotheses about RBP function, as enrichment for the ribosomal RNA precursor corresponds with RBPs regulating ribosomal RNA maturation whereas enrichment for retrotransposable elements corresponds to both regulation of retrotransposition itself as well as suppression of improper RNA processing due to cryptic elements contained within these elements. Binding maps across meta-profiles of mRNAs and exon-intron junctions similarly shows that RBP binding patterns correlate with RBP functional roles, and analysis of spliceosomal components indicates that eCLIP can be used to identify branch points and provides evidence for a 3’ splice site scanning model. In summary, these results provide further validation of the power of integrated analyses of RBP target maps generated by eCLIP in identifying novel principles of RNA biology, as well as generating RBP-specific hypothesis for further functional validation.

## Results

### Large-scale profiling of RNA binding protein binding sites with eCLIP

The eCLIP methodology enabled highly efficient identification of RBP binding sites [18], leading to the generation of the first large-scale database of RNA binding protein targets profiled in the same cell-types using a standardized workflow [20]. This dataset contains 223 eCLIP profiles of RNA binding sites for 150 RNA binding proteins (120 in K562 and 103 in HepG2 cells), covering a wide range of RBP functions, subcellular localizations, and predicted RNA binding domains (Fig. 1a; Supplemental Table 1-2). Each experiment contains biological duplicate immunoprecipitation libraries along with a paired size-matched input from one of the two experimental biosamples (Fig. 1b). For each experiment, raw sequencing data, processed data (including read mapping and identified binding sites), and experimental metadata (including antibody and immunoprecipitation validation documentation, biosample information, and additional related ENCODE datasets) were deposited at the ENCODE data coordination center (https://www.encodeproject.org) [20].

**Fig. 1.**
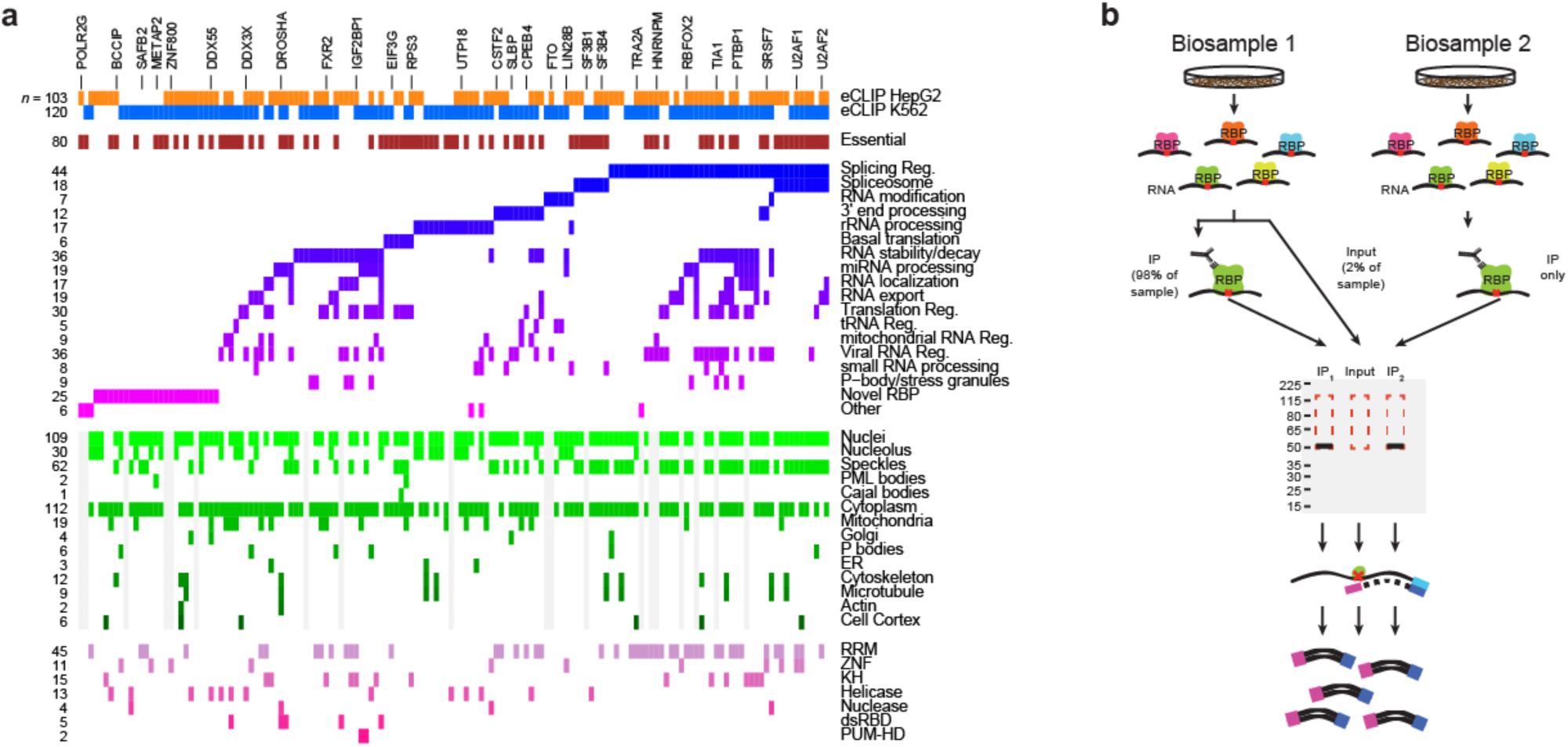
223 eCLIP datasets profile targets for 150 RNA binding proteins. (a) Colors indicate RBPs profiled by eCLIP, with manually annotated RBP functions, subcellular localization patterns from immunofluorescence imaging, and predicted RNA binding domains indicated (Supplemental Table 2). (b) Schematic overview of eCLIP as performed in the datasets described here. Two biological replicates (defined as biosamples from separate cell thaws and crosslinked more than a week apart) were performed for each RBP, along with one size-matched input taken from one of the two biosamples prior to immunoprecipitation.

Many CLIP methods included radioactive labeling of the 5’ end of RNA fragments with ^32^P to visualize protein-RNA complexes after SDS-PAGE electrophoresis and membrane transfer in order to query whether RNA bound to co-purified RBPs of different size is present [4]. However, the eCLIP protocol we utilized above did not include this direct visualization of protein-associated RNA due to the complexity of incorporating radioactive labeling at this scale, preferring validation of eCLIP signal with orthogonal approaches (such as comparison with *in vitro*-derived motifs or overlap with knockdown/RNA-seq changes). However, to address this question for future large-scale eCLIP profiling, we pursued alternative labeling approaches. We found that ligation of biotinylated cytidine (instead of the normal RNA adapter) enabled visualization similar to that observed with ^32^P while using commercially available chemiluminescent detection reagents for biotin-labeled nucleic acids (Sup. Fig. 1a-c). We note that unlike ^32^P labeling (which is done as a 5’ phosphorylation reaction with T4 Polynucleotide Kinase), this labeling uses the standard eCLIP RNA adapter ligation reaction and thus may more accurately reflect true protein-coupled RNA positioning.

Surprisingly, when expanding this approach across RBPs, we observed detectable transfer of RNA from non-crosslinked cells to nitrocellulose membranes in a supplier-dependent manner (Sup. Fig. 1b). We had previously noted that certain sourced nitrocellulose membranes contained greater amounts of RNA, which would then be recovered during library preparation (particularly in input libraries, which lack adapter addition prior to membrane transfer) [21]. However, we now observed that the recommended (lower contaminant, membrane I) membrane from that effort showed increased transfer of RNA than our previous supplier (membrane G) (Sup. Fig. 1d-f). Although the signal observed in crosslinked samples was typically significantly higher (median 12.5-fold across 17 RBPs tested), with 88% (15 out of 17) RBPs greater than 5-fold (Sup. Fig. 1d), for 2 out of 17 we observed within 5-fold RNA transfer in non-crosslinked samples (Sup. Fig. 1d,f).

To directly query whether this led to artifactual eCLIP peak identification, we chose seven eCLIP experiments performed with membrane I and performed replicate experiments with membrane G. Using MATR3 as an example, we observed that peak fold-enrichment compared across membranes was similar to that observed for within-membrane replicates (Sup. Fig. 1g). Extending this to all seven RBPs, only one (FXR2) out of seven showed notably lower replication of peak significance using membrane G (Sup. Fig. 1h), and even in that case we observed high overall correlation in peak fold-enrichment (Sup. Fig. 1i). Conservation of signal was not limited to peak calls, as we observed similar enrichments for retrotransposable and other RNA elements as well (Sup. Fig. 1j). Thus, although our data indicates that whether RNA that is not crosslinked to protein will transfer to nitrocellulose membranes is supplier- and product-dependent, but that it does not generally appear to add significant background to the eCLIP profiles studied here.

### Recovering RNA binding protein association to retrotransposons and other multicopy RNAs

Standard peak analysis revealed a wide variety of binding modes to mRNAs, with RBPs enriched for coding sequences, 3’ and 5’ untranslated regions, proximal and distal intronic regions, and non-coding RNAs [20]. Notably, we observed that RNA binding protein mRNAs were 1.4-fold enriched (*p* = 2.1×10^-22^ by one-sample t-test) among all peak-containing genes (median 13.5% per dataset, relative to 9.4% of all genes with at least one peak). In particular, well-studied splicing regulators (e.g. SRSF7 and TRA2A) were more than 3-fold enriched for binding to RBPs (Sup. Fig. 2a-b). In contrast, transcription factors were unchanged (1.0-fold depleted), suggesting that RNA processing regulators are particularly likely to themselves be the target of RNA processing regulation. In total, RBPs profiled in this study bound a median of 107 RBPs and 34 transcription factors, confirming the presence of a highly complex regulatory network of RNA and DNA processing (Sup. Fig. 2a).

In addition to single-copy RNA transcripts, the human genome contains many high-copy regions that are expressed as functional RNAs but present a substantial challenge to standard short read mapping strategies. These include RNAs such as the large and small ribosomal RNA (rRNA), 7SK snRNA, and others that have one or few expressed primary transcripts but dozens to hundreds of pseudogenes throughout the genome, as well as retrotransposable elements including LINE and Alu elements with thousands of moderately divergent sense and antisense copies throughout transcribed genes [22]. We found that simply including non-uniquely-mapped reads in standard analysis created thousands of peaks in introns, intergenic regions, and at pseudogenes that typically lacked standard peak shapes (likely reflecting sequencing errors relative to the main expressed transcript), indicating the need for improved methods to properly quantify RBP binding to such loci.

In order to include these RNA types in eCLIP analysis, we developed a ‘family-aware mapping’ approach in which adapter-trimmed reads are first mapped against a database of sequences for primary transcripts and pseudogenes for 36 families (Fig. 2a) (Supplemental Table 3). Reads mapping to reference transcripts contained within a family (e.g. LINE, YRNA, or 18S rRNA) are used for quantitation, but reads that map to multiple families are masked (discarding an average of 1.1% of reads). These results are then integrated with standard unique genomic mapping, incorporating reads that uniquely map to regions annotated as repetitive elements by RepeatMasker [23] into the final family quantitation (Fig. 2a). Confirming the success of this approach, we observed that in eCLIP replicates of YRNA-associating factor TROVE2/RO60 in K562 only 3.7 and 6.8% (replicate 1 and 2 respectively) of usable reads uniquely mapped to YRNA transcripts with standard processing (2.9 and 5.1% to RNY1/2/4/5, with another 0.7% and 1.8% to YRNA pseudogenes) (Fig. 2b). In contrast, for these same datasets 14.2% and 21.7% of reads mapped uniquely to the YRNA family using the family-aware mapping approach, making use of hundreds of thousands of additional reads that did not uniquely map to individual transcripts (Fig. 2b). Performing this analysis for all RBPs, we observed a wide range of read recovery and enrichment for particular elements (Fig. 2c, Supplemental Table 4). For some RBPs such as RPS11 (K562) an average of 95.2% of reads were only recovered using family mapping (68.1% mapping to RNA18S with an additional 24.1% to RNA28S). In contrast, only 10.4% of reads in KHSRP (K562) eCLIP mapped to multicopy family elements, with 58.9% uniquely mapping to the genome (including 41.1% uniquely mapping to introns outside of RepeatMasker elements) (Fig. 2c).

**Fig. 2.**
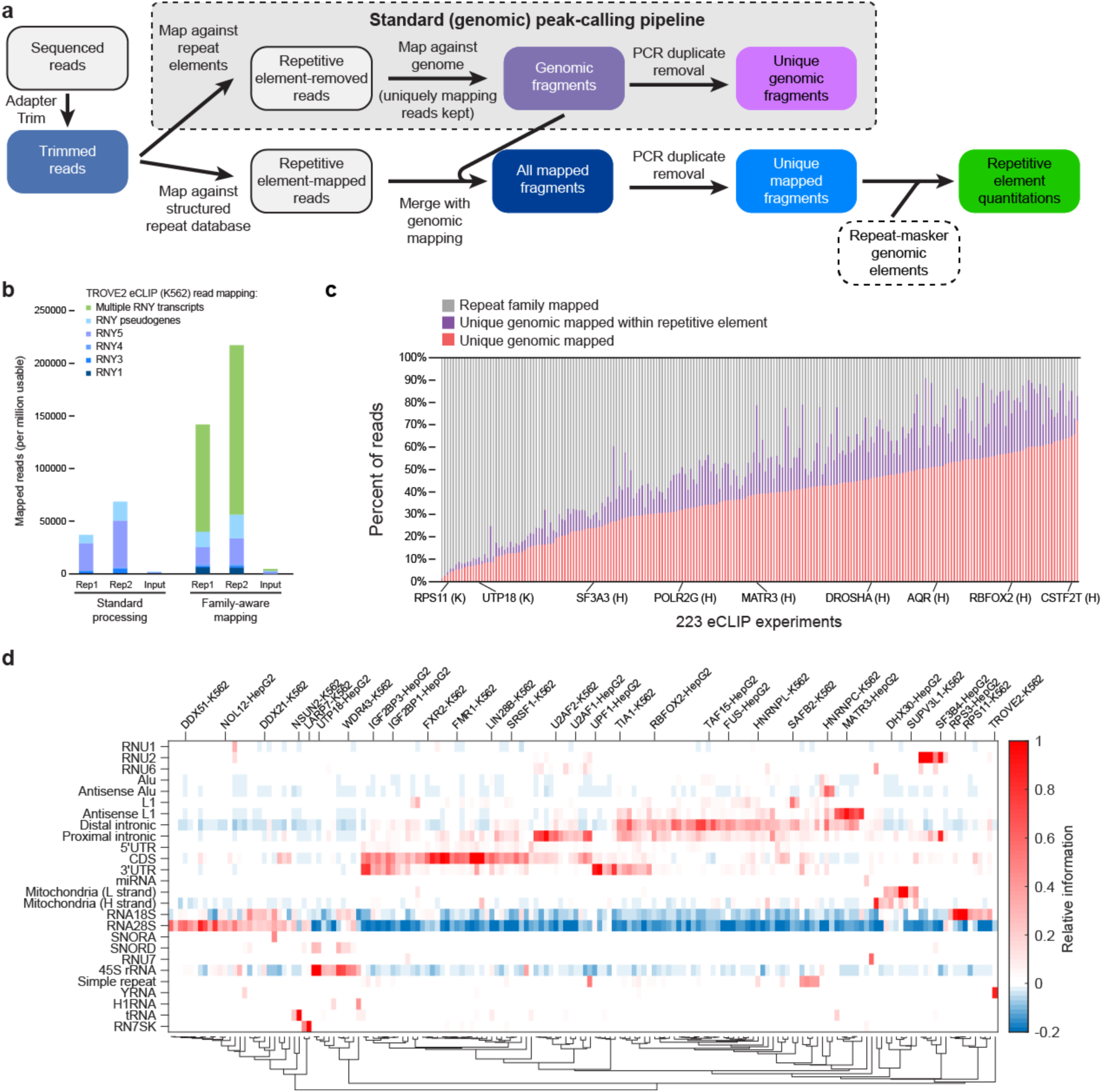
Quantification of repetitive elements and other non-uniquely mapped reads. (a) Schematic indicates major steps for (grey box) the standard eCLIP peak calling pipeline (which uses uniquely mapping reads only) and the repetitive element quantification pipeline. (b) Stacked bars indicate the number of reads from TROVE2 eCLIP in K562 that map either uniquely to one of four primary Y RNA transcripts, map uniquely to Y RNA pseudogenes (identified by RepeatMasker), or (for family-aware mapping) map to multiple Y RNA transcripts but not uniquely to the genome or to other repetitive element families. (c) Stacked bars indicate the fraction of reads (averaged between replicates) of all 223 eCLIP experiments, separated by whether they map (red) uniquely to the genome, (purple) uniquely to the genome but within a repetitive element identified by RepeatMasker, or (grey) to repetitive element families. Datasets are sorted by the fraction of unique genomic reads. (d) Heatmap indicates the relative information for 26 elements and 168 eCLIP datasets, requiring elements and datasets to have at least one entry meeting a 0.2 relative information cutoff (based on Sup. Fig. 2d). See Table 1 for RBP:element enrichments meeting this criteria.

**Table 1.**
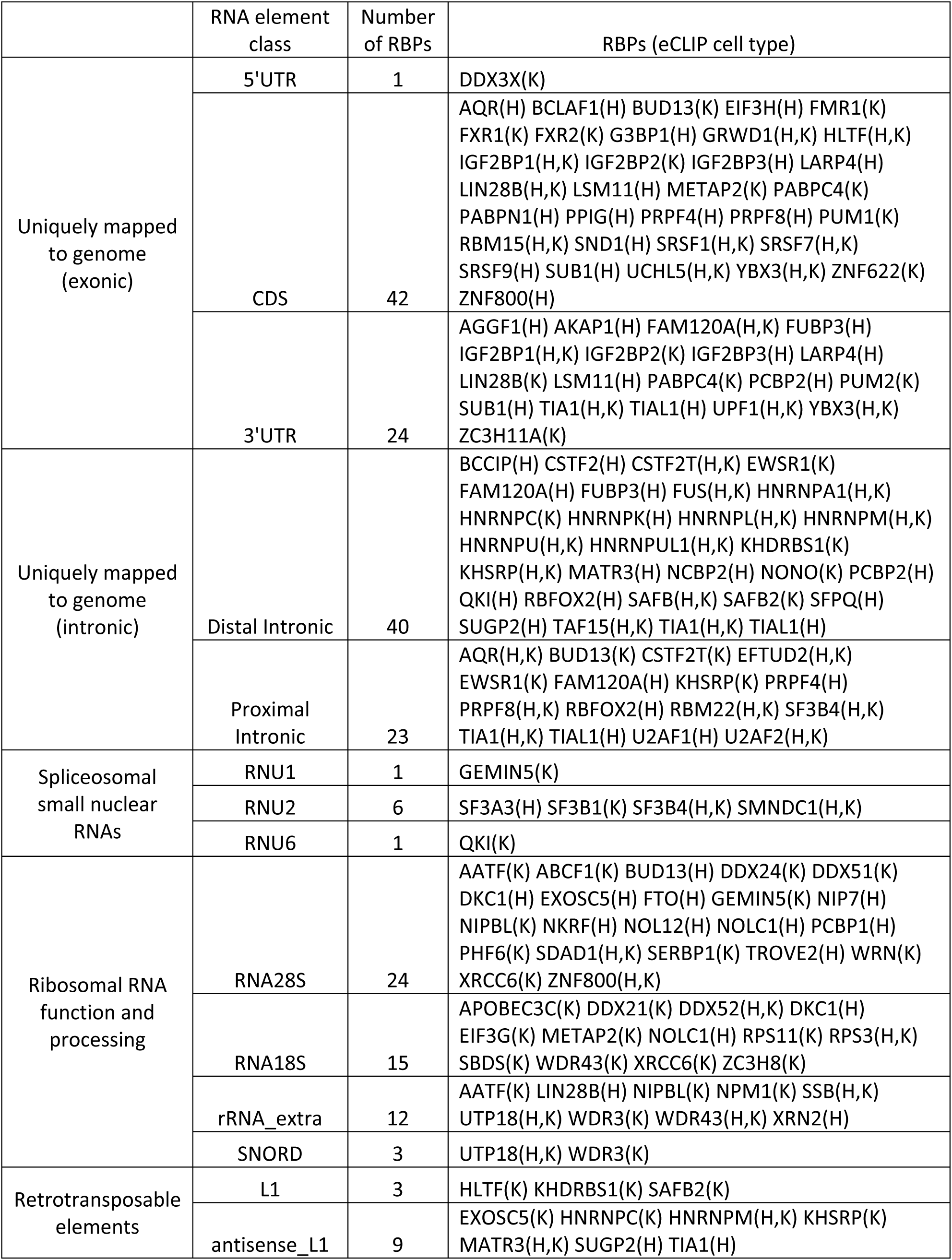
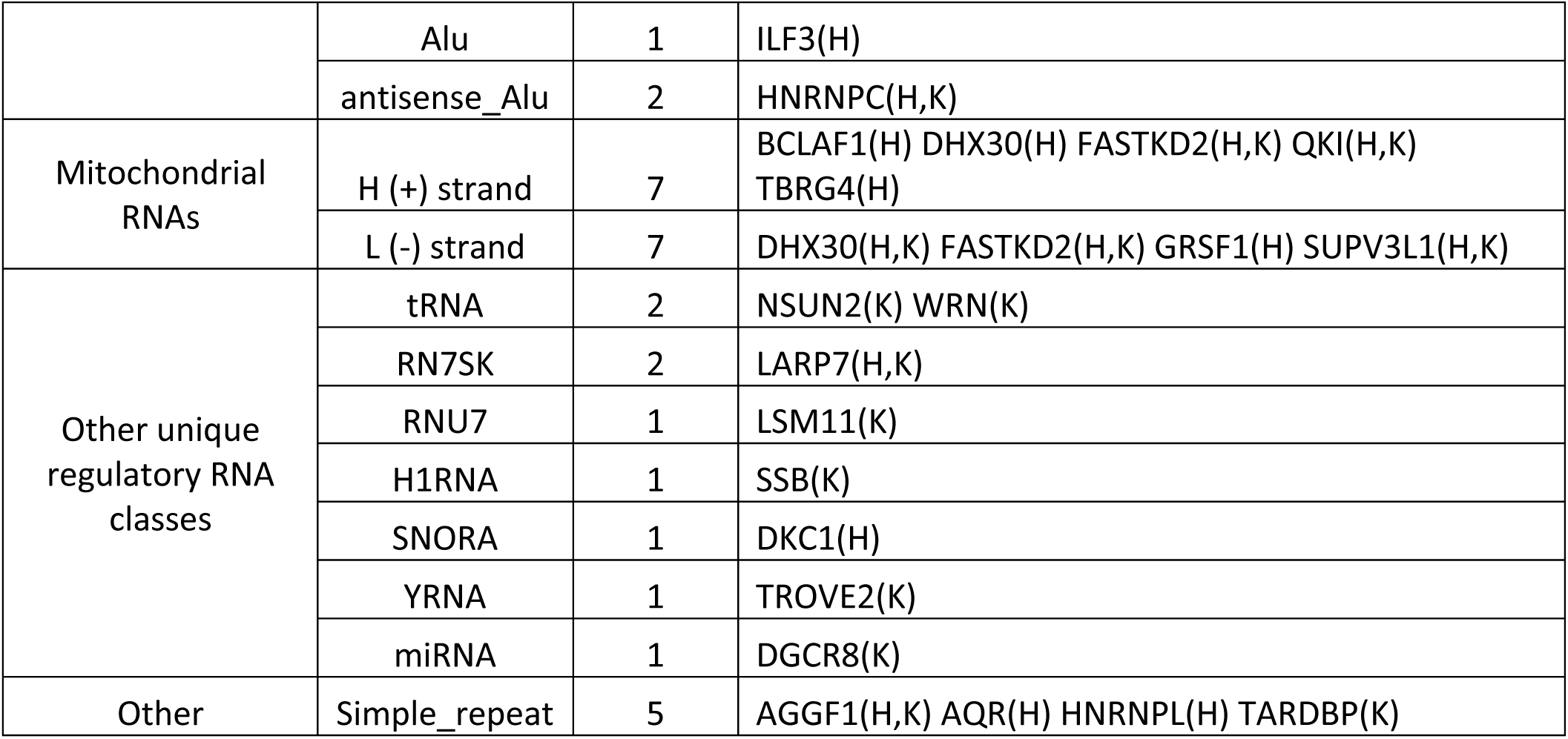
Major RNA element families enriched in RNA binding protein eCLIP experiments.

At the element-level, our family-aware mapping strategy recovers many known processing or interacting factors, including RBPs enriched for the mature 18S (RPS3, RPS11) and 28S rRNA (DDX21, NOL12) as well as the 45S rRNA precursor (UTP18, WDR43), tRNAs (NSUN2), RN7SK (LARP7), YRNA (TROVE2) and others (Fig. 2d). To validate this approach, we considered 17 RNA elements with well-studied direct links to either RBP function (such as snoRNA binding with rRNA processing and snRNA binding with snRNA processing and the spliceosome) or specific RBP regulators (e.g. snRNA RN7SK with LARP7 [24] and YRNAs with TROVE2/Ro60 [25]) (Sup. Fig. 2c). We observed that 140 eCLIP datasets had one of these 17 elements as the most highly enriched (by relative information, which we observed to better enable comparison across elements versus fold-enrichment), and in 84 (60%) of these cases the RBP was previously characterized as having the element-paired RBP function, indicating that this approach is highly successful at recovering targets that reflect annotated functions of profiled RBPs. To set a cutoff for analysis, we found that an information cutoff of 0.2 maximized predictive accuracy, at which 70% (74 out of 105 RBPs with the most enriched RNA element meeting this cutoff) had annotated functions matching the known role for this element (Sup. Fig. 2d). Using this cutoff, 235 RBP-element pairings were identified with large numbers of RBPs associated with mRNA regions (42 with CDS, 24 with 3’UTR, 40 with distal intronic, and 23 with proximal intronic regions) and rRNA (24 with RNA28S and 15 with RNA18s, as well as 12 with precursor 45S rRNA), and smaller numbers associated with other specific RNA classes (Figure 2d, Table 1).

### Characterization of ribosomal RNA interactors and processing factors

Ribosomal RNA (rRNA) is the most abundant RNA found in eukaryotic cells and plays essential roles in defining the structure and activity of the ribosome. In humans, the 5S rRNA is separately transcribed, whereas the 18S, 28S, and 5.8S rRNAs are transcribed as one 45S precursor transcript that then undergoes a complex series of cleavage and RNA modification steps to process the mature rRNAs, which then form complex structures that scaffold the assembly of ∼80 proteins to create the functional ribosome [26]. Unbiased approaches have characterized over 250 additional factors as playing critical roles in processing pre-rRNA, indicating that rRNA processing and function represents a major function of RBPs in humans [27].

Considering the 150 RBPs profiled, we observed that different subsets of RBPs showed enrichment to specific rRNAs (Fig. 3a), suggesting that the incorporation of normalization against paired input was successful in removing general background at abundant transcripts. Although we are unable to distinguish between mapping to mature 18, 28, and 5.8S transcripts versus those regions in the precursor, the ∼4-fold lower read density we observe for 45S (median 433 reads per million (RPM)) versus 18S (2371 RPM) or 28S (1794 RPM) in eCLIP input samples (Sup. Fig. 3a-c) suggests that the majority of 18S and 28S reads reflect mature rRNA transcripts. Considering 36 RBPs previously shown to effect pre-rRNA processing [27], we found that 16 (44.4%) had rRNA processing-related elements as the most enriched (at a 0.064 position-wise information cutoff) relative to 12.3% of others (3.6-fold enriched, p = 8.4×10^-5^ by Fisher’s Exact test) (Sup. Fig. 3d). Despite high and relatively even read density overall on the abundant rRNA transcripts (Sup. Fig. 3a-c), we observed that these rRNA-enriched RBPs showed a number of specific enrichment patterns: two on the 45S precursor (one situated around the 01 and A0 early processing sites, and a second located ∼2000nt further downstream that is discussed below), a cluster at position ∼4200 of the 28S, and a cluster at ∼1150 of the 18S, along with other profiles unique to individual RBPs (Fig. 3a). Distinct ribosomal components RPS3 and RPS11 had different positional enrichments, as expected given their different positioning within the 18S ribosome (Sup. Fig. 3e).

**Fig. 3.**
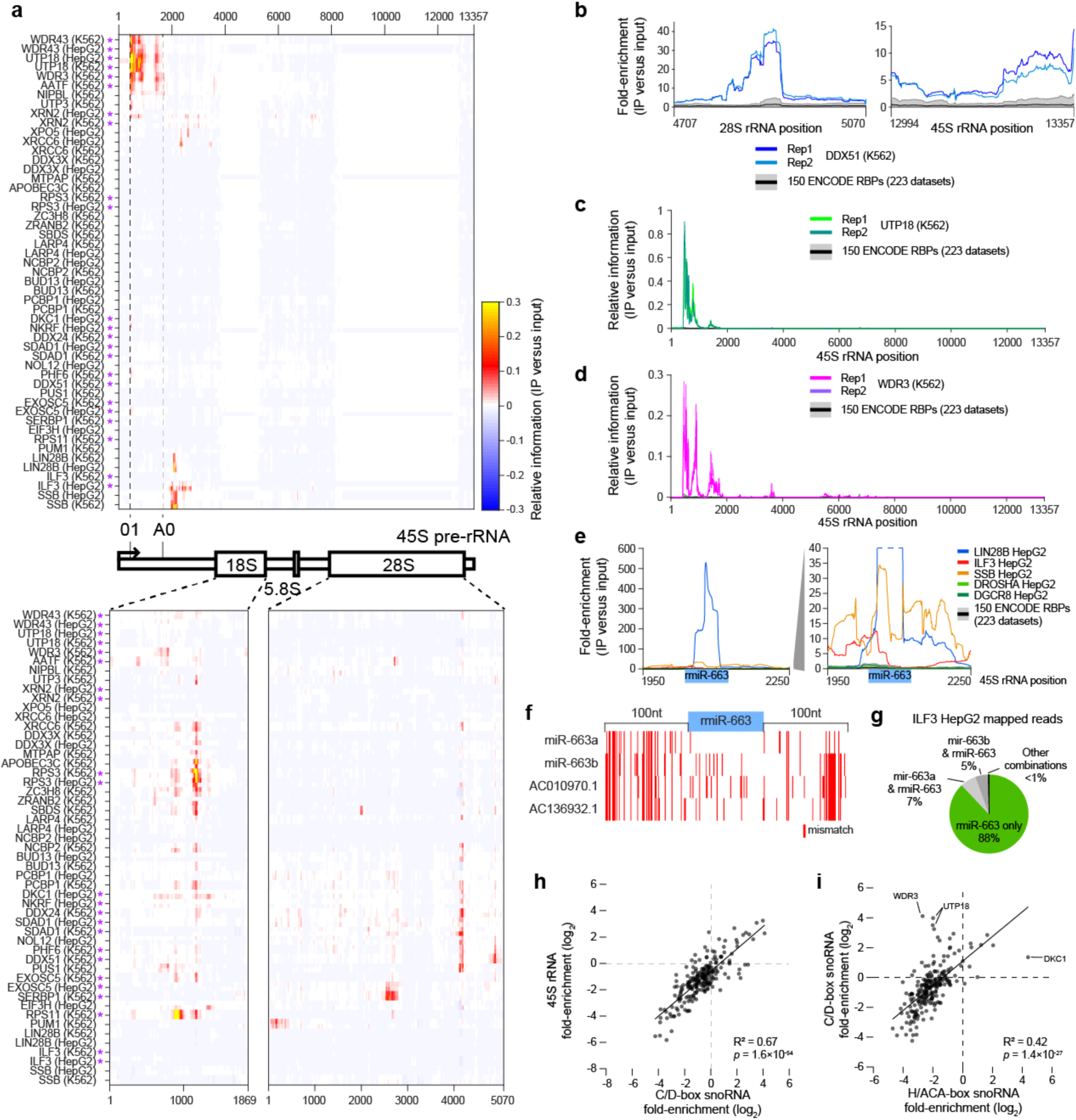
eCLIP enrichment for rRNA links RBPs with ribosomal RNA processing. (a) Heatmap indicates relative information at each position along (top) the ribosomal RNA precursor 45S polycistronic transcript and (bottom) within the mature 18S and 28S transcripts. Reads mapping equally to the 45S and mature 18S or 28S are assigned to the mature for quantitation. Purple asterisk indicates RBPs for which knockdown showed rRNA processing defects in Tafforeau *et al*. [27]. (b) Lines indicate fold-enrichment in DDX51 eCLIP in K562 cells at the 3’ end of the 28S and 45S transcript. For this and further plots, black line indicates mean and grey region indicates 10^th^ to 90^th^ percentile across all 223 eCLIP datasets. (c-d) Lines indicate relative information for (c) UTP18 in K562 and (d) WDR3 in K562 across the 45S precursor. (e) Lines indicate fold-enrichment for indicated RBPs within a region flanking putative ribosomal-encoded microRNA rmiR-663. (f) Red indicates mismatch positions relative to ribosomal rmiR-663 (and 100nt flanking regions) for genomic-encoded miR-663a, miR-663b, and two additional homologous regions containing putative microRNAs. (g) Pie chart indicates the fraction of reads in ILF3 HepG2 eCLIP mapping (green) with fewer mismatches to rmiR-663, or (grey) mapping equally well to rmiR-663 and other miR-663 family members as indicated. See Supplemental Fig. 3j-k for LIN28B (HepG2) and SSB (HepG2). (h-i) Points indicate fold-enrichment in each eCLIP dataset for (h) C/D-box snoRNAs versus 45S precursor RNA, and (i) H/ACA-box snoRNAs versus C/D-box snoRNAs. Pearson correlation and significance was calculated in MATLAB.

Our data on rRNA precursor position-specific enrichment confirms and provides further resolution to proteins previously characterized to play roles in ribosomal RNA processing. Some factors had specific positioning, including DDX51 which had specific enrichment at the 3’ end of 28S as well as the 3’-ETS precursor region, consistent with previous characterization of the role of DDX51 in 3’ end maturation of 28S [28], and UTP18 which had specific enrichment at the 5’ end, matching its roles in early cleavages at the 01, A0, and 1 sites suggested from large-scale screening data [27] (Fig. 3b-c, Sup. Fig. 3f-g). Others, such as WDR3, had broader enrichment patterns that suggest participation in multiple maturation steps (Fig 3d, Sup. Fig. 3h).

Surprisingly, we observe a cluster of RBP association in the 45S precursor around position 2100, a region located between the A0 and 1 processing sites which lacks a well-defined processing role (Fig. 3a) [26]. Two of these factors have previous links to nucleolar activity, as ILF3 (also known as NF90) was previously shown to associate with pre-60S ribosomal particles in the nucleolus and knockdown of ILF3 gives defects in rRNA biogenesis [27, 29], and LIN28B has been shown to repress let-7 processing by sequestering pri-let-7 in the nucleolus [30]. In this region, multiple sites of ILF3 and SSB enrichment flank a more specific region enriched in LIN28B eCLIP (Fig 3e, Sup. Fig. 3i) which has previously been described to contain a potential rRNA-encoded microRNA, rmiR-663a [31]. As rmiR-663a shares similar sequence to genomic-encoded miR-663a on chromosome 20 (and would have the same mature miRNA sequence), it has been challenging to isolate expression of the ribosomal-encoded transcript in isolation [32], and indeed the majority of LIN28B eCLIP reads mapping to pri-miRNA map equally to both variants (Sup Fig. 3j). However, when we used sequence variants in the pri-miR sequence as well as the more variable flanking sequence to estimate their separate expression (Fig. 3f), we observed that reads unique to the rmiR outnumbered those unique to genomic homologs by more than 400-fold (Fig. 3g & Sup. Fig. 3j-k), indicating that the observed signal is likely derived from 45S rather than other genomic homologs.

Finally, we considered binding to snoRNAs, a class of highly structured small RNAs that play essential roles in guiding modification of ribosomal RNAs. We found that enrichment for C/D-box snoRNAs, which canonically guide methylation of RNA, was highly correlated to enrichment for the 45S precursor (R^2^ = 0.67, p = 1.6×10^-54^)(Fig. 3h), providing further confirmation that these 45S-enriched RBPs are likely playing key roles in rRNA processing. Surprisingly, however, we observed that enrichment for H/ACA-box snoRNAs showed far lower correlation with enrichment for either C/D-box snoRNAs (R^2^ = 0.42) or the 45S precursor (R^2^ = 0.17) (Fig. 3i, Sup. Fig. 3l). Thus, this data confirms the ability of eCLIP with input normalization to specifically isolate enrichment between abundant snoRNA classes, and suggests that (at least for the RBPs profiled to date here) we see stronger overlap between rRNA precursor and C/D-box versus H/ACA-box snoRNAs.

### Repetitive elements define a significant fraction of the RBP target landscape

Repetitive elements constitute a large fraction of the non-coding genome [33], and elements annotated by RepBase constitute an average of 12.2% of reads observed in eCLIP input experiments (Sup. Fig. 4a). In particular, as retrotransposable L1/LINE and Alu elements constitute 10.8% and 0.4% of intronic sequences respectively (Sup. Fig. 4b), they represent a significant fraction of the pool of nuclear transcribed pre-mRNAs available for RBP interactions. Although some RBPs have been shown to play roles in regulation of active retrotransposition [34], the majority of intronic elements have accumulated mutations or deletions and are no longer capable of active retrotransposition, leaving the question of their function relatively poorly understood. However, recent analyses of RBP targets identified by CLIP (including early releases of the eCLIP data considered here) have shown that both antisense Alu and antisense LINE elements contain cryptic splice sites that can lead to improper splicing and polyadenylation, suggesting that a major yet unappreciated role for many RBPs may be to suppress the emergence of inappropriate cryptic RNA processing sites introduced upon retrotransposition [35, 36].

Querying for RBPs with enriched eCLIP signal at retrotransposable and other repetitive elements, we surprisingly observed that only a small subset of elements (notably including L1 and Alu elements both in sense and antisense orientation) showed high RBP specificity, whereas most elements showed extremely highly correlated enrichments across RBPs (Fig. 4a, Sup. Fig. 4c). This group of elements showed enrichment in a small subset of eCLIP experiments, notably including multiple members of the highly abundant HNRNP family (HNRNPA1, HNRNPU, HNRNPC, and HNNRPL), indicating that they may be coordinately regulated to prevent inappropriate RNA processing.

**Fig. 4.**
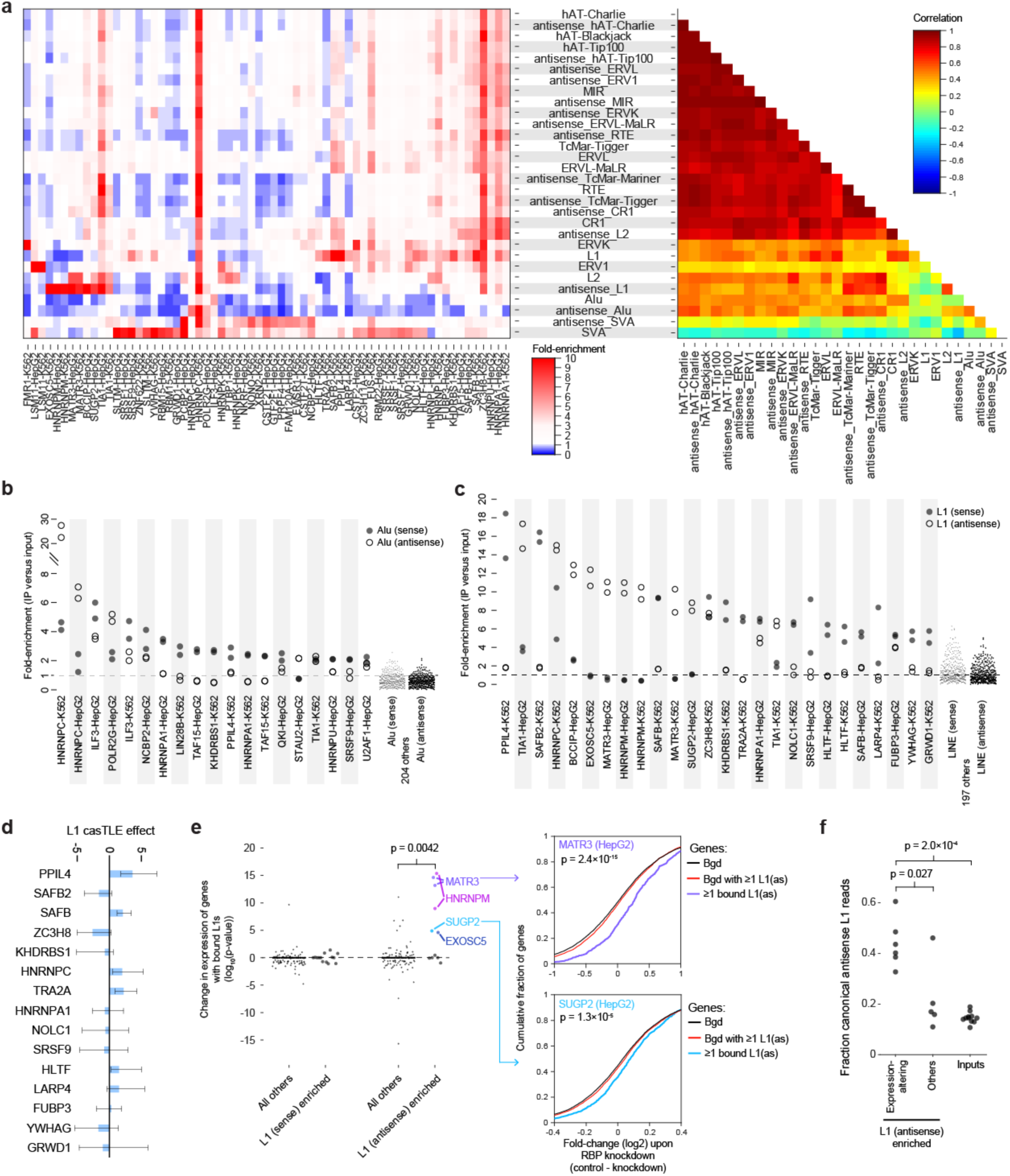
RBP association at retrotransposable and other repetitive elements. (a) (left) Heatmap indicates fold-enrichment in eCLIP versus paired input, averaged across two biological replicates. Shown are 30 RepBase elements which had average RPM>100 in input experiments and at least one RBP with greater than 5-fold enrichment and 65 eCLIP experiments with greater than 5-fold enrichment for at least one element. (right) Color indicates correlation in fold-enrichment between elements across the 65 experiments. (b-c) Points indicate fold-enrichment for (b) Alu elements and (c) L1 LINE elements in individual biological replicates. Shown are all RBPs with average enrichment of at least 2 (for Alu elements) or 5 (for L1 elements). (d) Bars indicate L1 retrotransposition casTLE effect score (positive score indicates increased retrotransposition upon RBP knockout), with error bars indicating 95% minimum and maximum credible interval estimates (data from Liu *et al*. [39]). (e) (left) Each point indicates significance (from two-sided Kolmogorov-Smirnov test) between fold-changes observed in RNA-seq of RBP knockdown for the set of genes with one or more RBP-bound L1 (or antisense L1) elements versus the set of genes containing one or more L1 (or antisense L1) elements but lacking RBP binding (defined as overlap with an IDR peak). RBPs were separated based on requiring 5-fold enrichment for L1 elements as in (c). (right) Cumulative distribution plots for (top) MATR3 in HepG2 and (bottom) SUGP2 in HepG2. Significance shown is versus the set of genes containing one or more L1 (or antisense L1) elements but lacking RBP binding (red line). (f) Points indicate the fraction of antisense L1-assigned reads that map to canonical (RepBase) elements for six expression-altering antisense L1-enriched eCLIP datasets (from (e)), five other antisense-L1 enriched eCLIP datasets, and 11 paired input samples. Significance is from two-sided non-parametric Kolmogorov-Smirnov test. See Sup. Fig. 4g for full distribution of read assignments.

Analysis of Alu elements recapitulated a previously described interaction of HNRNPC with antisense Alu elements [35], but additionally revealed two RBPs with more than 5-fold enrichment: ILF3 (enriched for both sense and antisense Alu elements) and RNA Polymerase II component POLR2G (antisense) (Fig. 4b, Sup. Fig. 4d). Both of these factors have previous links to RNA processing through Alu elements, as ILF3 association at was suggested to repress RNA editing in Alu elements [37] and Alu elements have been shown to effect RNA Polymerase II elongation rates [38]. In total, 19 datasets showed more than 2-fold enrichment for either Alu or antisense Alu elements (Fig. 4b).

Considering L1/LINE elements, we observed enrichment with far more RBPs, with 26 datasets showing 5-fold enrichment (Fig. 4c). Interestingly, we observed generally distinct sets for sense versus antisense L1 enrichment, with only HNRNPC (in K562, but not HepG2) and ZC3H8 showing enrichment for both (Fig. 4c, Sup. Fig. 4e). The RBPs identified here align well with those identified in an independent analysis of L1-associated RBPs which used a subset of these datasets along with independent iCLIP and other datasets, confirming robustness of this analysis across different approaches to quantify enrichment to L1 elements [36]. To query the role of L1 association, we first considered whether binding could specifically act to repress L1 retrotransposition itself. Of the 15 RBPs with more than 5-fold enrichment at sense L1 elements, SAFB (p = 0.002), PPIL4 (0.06) and TRA2A (p = 0.05) were all identified as candidate suppressors of L1 retrotransposition in a recent genome-wide CRISPR screening assay [39], suggesting that this eCLIP enrichment approach identifies functional regulators of retrotransposition (Fig. 4d).

However, we observed that while enriched signal was centered at L1 sense and antisense elements, the signal often extended for multiple kilobases on either side (Sup. Fig. 4f), indicating that despite the overlap with functional regulators of active lines, the majority of eCLIP signal is likely coming from inactive L1 elements contained within pre-mRNAs rather than independently transcribed active L1 elements in the cell lines studied here. Thus, we next assayed whether these RBPs showed evidence for silencing cryptic RNA processing sites created upon retrotransposition, as previously described [35, 36]. To do this, we hypothesized that knockdown of such RBPs would lead to inclusion of premature stop codons that signal nonsense mediated decay, ultimately decreasing abundance of target mRNA transcripts. For MATR3, we indeed observed that genes containing one or more antisense L1 elements overlapped by peaks showed significantly decreased expression upon RBP knockdown (Fig. 4e), consistent with recent findings that MATR3 binding blocks both cryptic poly(A)-sites and splice sites within LINEs [36]. Interestingly, we observed a similar pattern for 3 other RBPs with antisense L1 enrichment, HNRNPM (which has been identified in complexes with MATR3 [40]), SUGP2, and EXOSC5 (Fig. 4e). These four RBPs also showed particular enrichment for reference L1 sequences as opposed to unique genomic mapping to more degenerate elements, suggesting that this specifically segregates expression-altering antisense L1-enriched RBPs (Fig. 4f, Sup. Fig. 4g).

### Metagene binding profiles reveal RBP functions

Next, we turned to the question of whether eCLIP peak distributions could reveal RBP roles in mRNA processing. To better separate RBP association patterns, we considered the distribution peaks across a meta-gene generated by size-normalizing binding across all protein coding transcripts relative to transcription start and stop sites and start and stop codons, and then averaging across all expressed genes (Fig. 5a). Considering binding relative to the coding region (CDS) and 5’ and 3’ untranslated regions of spliced mRNA, we observed an overall average of approximately one peak per gene across the entire mRNA (Sup. Fig. 5a), with a variety of patterns of individual RBP association (Fig. 5b).

**Fig. 5.**
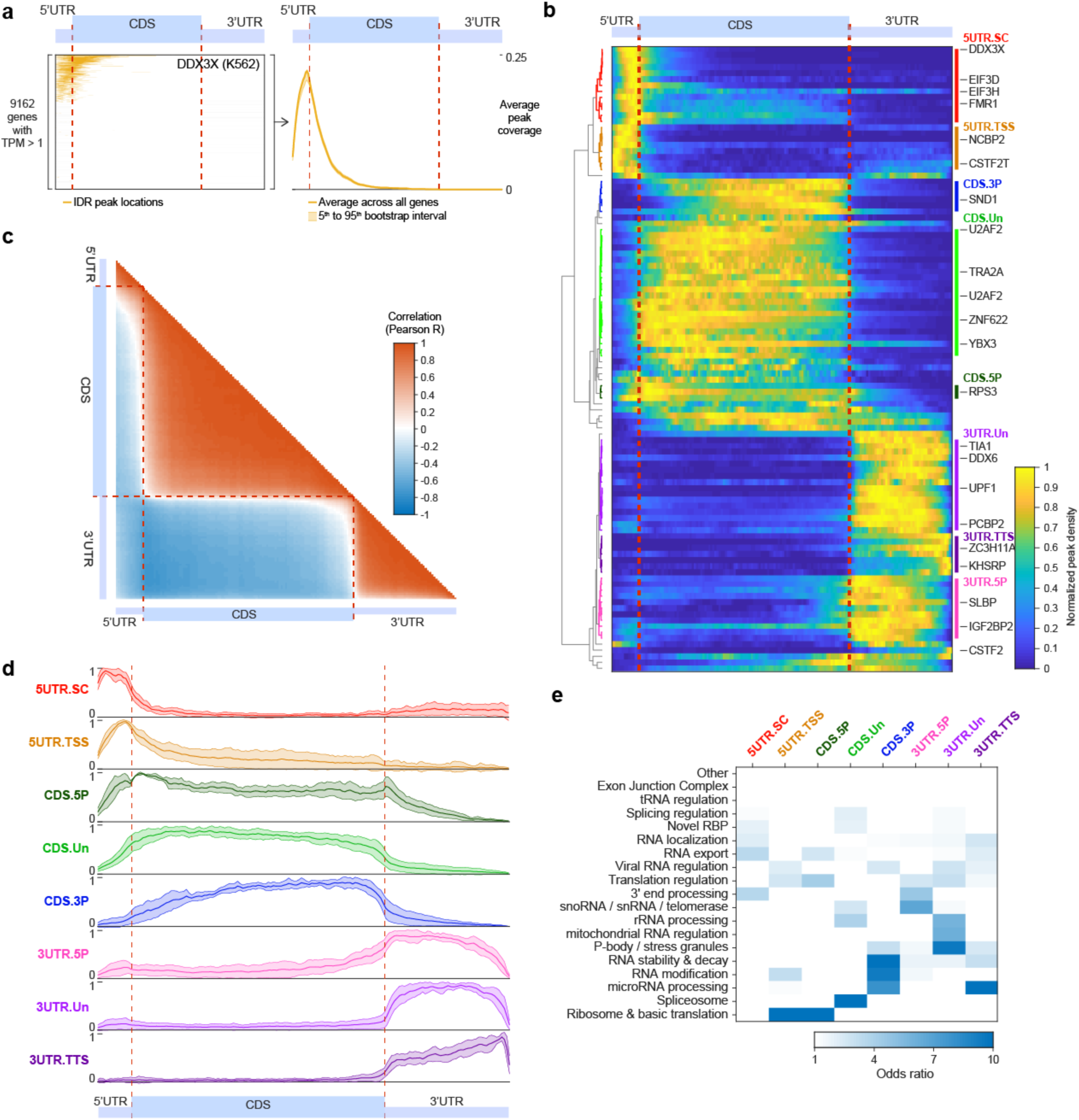
mRNA metagene profiles from eCLIP correspond to RBP regulatory roles. (a) (left) Each line indicates the presence (orange) of a reproducible DDX3X K562 eCLIP peak for 9,162 mRNAs that are expressed (TPM > 1) in K562. Each gene was normalized to 13 5’UTR, 100 CDS, and 49 3’UTR bins (based on average lengths among expressed transcripts in K562 cells). (right) A meta-mRNA plot is generated by averaging across all expressed genes, with shaded region indicating 5^th^ to 95^th^ percentile observed in 100 bootstrap samplings. (b) Heatmap indicates peak coverage for 104 datasets (requiring at least 100 reproducible peaks and at least one meta-mRNA position with 5^th^ percentile greater than 0.002), normalized by setting minimum value to zero and maximum to one. Meta-mRNA profiles were hierarchically clustered and manually labeled. (c) Heatmap indicates pair-wise correlation (Pearson R) between each pair of positions along the meta-mRNA in (b). (d) Lines indicate average normalized peaks per bin for all RBPs in the indicated class. Shaded region indicates one standard deviation. (e) Heatmap indicates odds ratio of overlap between eCLIP datasets in (x-axis) indicated meta-mRNA cluster versus (y-axis) annotated RBP functions. See Sup. Fig. 5d for significance.

At a global level, the most striking observation was clear delineation points at the start and stop codon positions (Fig. 5b-c), likely reflecting the fact that translation initiation is unique to the 5’UTR whereas the 3’UTR is the only region where bound RBPs will not be removed by translating ribosomes. However, more subtle clustering revealed distinct subgroups within the broader 5’UTR, CDS, and 3’UTR-enriched classes (Fig. 5b,d). For example, we observed two distinct classes of 5’UTR binding that appear to correlate with distinct RBP functions. The first (5UTR.TSS) showed greater enrichment closer to the transcription start site and included nuclear 5’ end processing factors such as cap-binding protein NCBP2 (Fig. 5b,d). In addition to 5’ end enrichment, this class also contained RBPs with substantial 3’UTR signal, such as 3’ end processing factor CSTF2T (which also showed significant signal extending past annotated transcription termination sites (Sup. Fig. 5b), consistent with previous CLIP studies [41]). A second set (5UTR.SC) showed biased peak presence closer to the start codon and included both canonical translational initiation factors (such as EIF3G, EIF3D, and EIF3H) as well as RBPs previously shown to play translational regulatory roles (including DDX3X, SRSF1, and FMR1) (Fig. 5b).

Similarly, we also observed distinctions within CDS binding, with either uniform (CDS.UN) density or biased towards the 5’ (CDS.5P) or 3’ (CDS.3P) end. We observed that 13 out of 15 spliceosomal RBPs showed CDS enrichment (10 of which fell into the CDS.UN category), likely reflecting the general lack of introns in 5’UTRs (due to their small size) and 3’UTRs (as they would create targets for nonsense-mediated decay) (Fig. 5b,d).

Finally, we observed multiple modalities of 3’UTR peak distribution. The 3UTR.Un class showed relatively uniform density and contained many well-characterized 3’UTR binding proteins, including NMD factor UPF1 and stress granule factor TIA1. In contrast, RBPs in the 3UTR.5P class had peak density enriched closer to (and continuing 5’ of) the stop codon, including the well-studied IGF2BP family of RBPs (Sup. Fig. 5c). Finally, we observed a number of RBPs with increased enrichment towards the transcription termination site (3UTR.TTS).

Next, we considered whether these patterns corresponded to different RNA processing functions. Although the number of RBPs is limited for some functions, we observed that many clusters had significant overlaps with distinct RBP functional annotations (Fig. 5e, Sup. Fig. 5d). In particular, RBPs associated with nuclear RNA processing steps showed little change (median 1.2-fold decrease in peak density around the stop codon, whereas RBPs with cytoplasmic roles showed a significant 1.6-fold increase (Sup. Fig. 5e), consistent with a stronger role for the stop codon as a delineation point for cytoplasmic RBP association. In all, our results suggest that the pattern of relative enrichment in different gene regions is predictive of the regulatory role that the RBPs play.

### Splicing regulatory roles revealed by intronic metagene profiles

Next, we performed regional analysis to query binding to exons (specifically 50nt bordering the splice sites) and 500nt of proximal introns flanking both the 3’ and 5’ splice sites. As an example, we observed that out of 89,265 introns present in highly expressed transcripts (TPM>1), 2,699 had a significant IDR peak from eCLIP of U2AF2 in K562 cells (Sup. Fig. 6a). These peaks had a stereotypical positioning at the 3’ splice site (extending into the downstream exon due to the use of full reads rather than just read 5’ ends for analysis), matching the well-characterized role of U2AF2 in 3’ splice site recognition (Fig. 6a). These matrices were then summed across all introns to calculate a meta-intron plot representing the average peak coverage at each position, with confidence intervals estimated by bootstrapping (Fig. 6b).

**Fig. 6.**
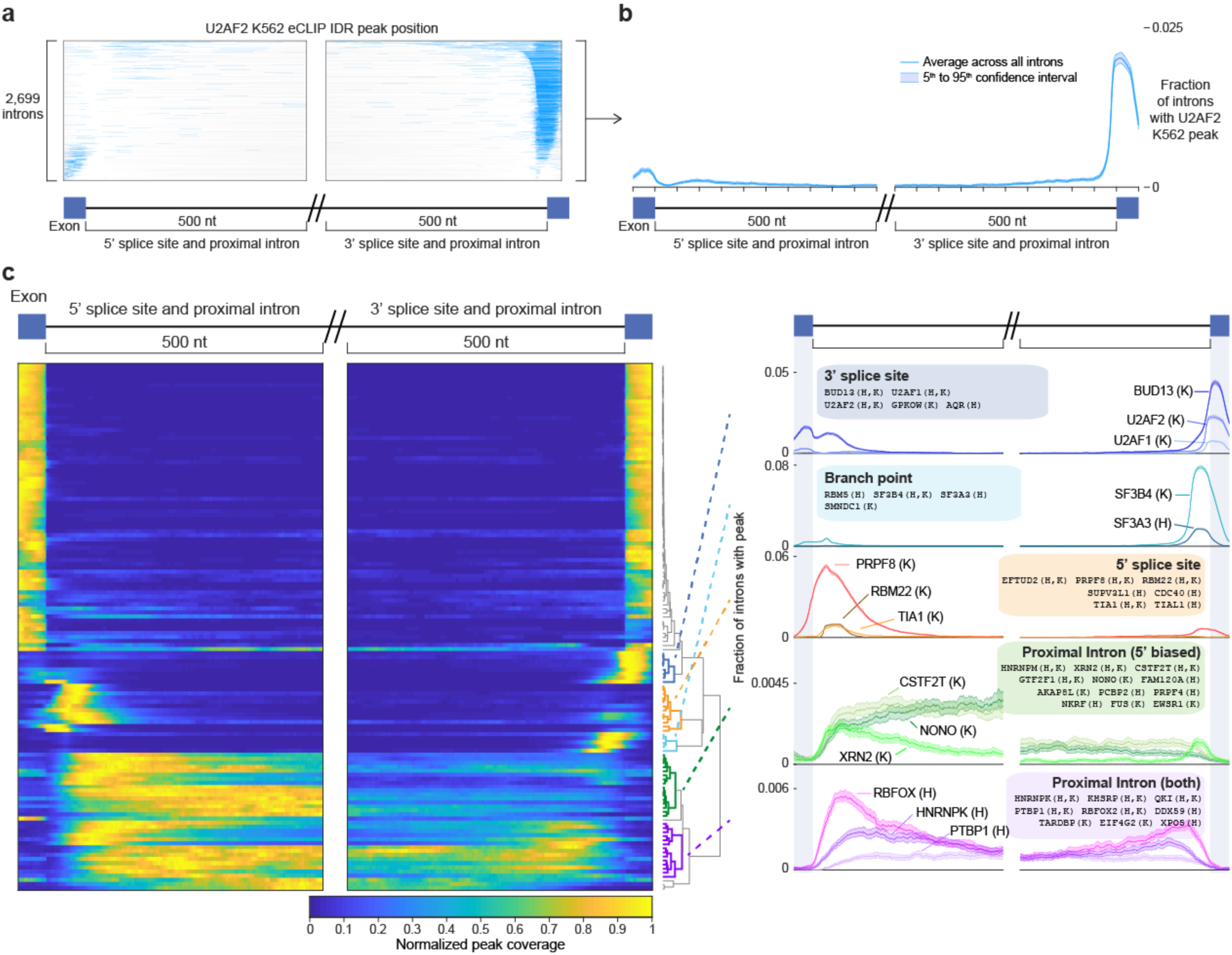
Meta-exon plots reveal intronic regulatory roles. (a) Each line indicates the presence (in blue) of a reproducible U2AF2 K562 eCLIP peak for 2,699 introns that contain at least one peak within the displayed region (500 nt of proximal intron and 50nt of exon flanking the 5’ and 3’ splice sites). See Sup. Fig. 6a for all 89,265 introns. (b) Meta-exon plot for data shown in (a), with line indicating average and shaded region indicating 5^th^ to 95^th^ percent confidence interval (derived by 100 bootstrap samplings). (c) (left) Heatmap indicates average peak coverage across all introns for 130 RBPs with at least 100 peaks and 5^th^ percentile confidence interval at least 0.0005 (for heatmap visualization, the maximum value for each dataset was set to one to calculate normalized coverage). (right) Lines show individual RBP examples for five clusters identified based on similar meta-exon profiles. Y-axis indicates fraction of introns with peak.

Performing this analysis for 130 RBPs with sufficient peaks (see Methods), we observed that the profiles recapitulated many known binding patterns, including U2AF1 and U2AF2 at the 3’ splice site, SF3B4 and SF3A3 at the branch point, PRPF8 at the 5’ splice site, and RBFOX2 and PTBP1 at proximal introns (Fig. 6c). Clustering analysis indicating a number of distinct RBP association patterns. In addition to a large group of exclusively exonic datasets, we observed clusters for the canonical splicing features (5’ splice site, 3’ splice site, and branch point), and two additional clusters: one where RBPs showed enrichment for peaks at proximal introns flanking both the 5’ and 3’ splice sites, and one with dominant enrichment in the 5’ splice site proximal intron only (Fig. 6c, right). We also observed a wide range of peak frequency; canonical splicing machinery components such as U2AF2, SF3B4 and PRPF8 had significantly enriched peaks at many introns (with a position maximum of 3.6%, 7.8%, and 5.3% of queried abundant introns respectively in K562), whereas factors such as PTBP1 and RBFOX2 were less commonly enriched at specific positions (0.1% and 0.5% respectively) (Fig. 6c).

### Insights into spliceosomal association and core splicing regulation

The breadth of RBPs profiled provided a unique opportunity to explore their interactions with the spliceosome and their impacts on splicing regulation. In addition to contacting the intron, many spliceosomal and splicing regulatory proteins also interact with the spliceosomal small nuclear RNAs (snRNAs). The overall snRNA family includes five specific RNA families (U1, U2, U4, U5, and U6, which also have variant isoforms that differ slightly in sequence) that play essential roles in canonical GT-AG RNA splicing, as well as four (U11, U12, U4atac, U5atac) specific to the minor AT-AC spliceosome, each of which plays specific mechanistic roles during splicing [42]. Thus, RBP association with a particular snRNA can help to map its function to a particular step in splicing. Quantitating snRNA enrichment using the family-aware mapping described above, we recapitulated many known associations between RBPs and the spliceosome, including interactions of SF3B4 with U2 snRNA (47- and 32-fold enriched in HepG2 and K562 respectively)[43] and GEMIN5 with U1 (11.2-fold enriched in K562)[44] (Fig. 7a). In some cases, these dominated overall RNA recovery; for example, an average of 41% of reads from SF3A3 eCLIP and 17% and 20% of SF3B4 eCLIP reads in HepG2 and K562 respectively mapped to the U2 snRNA, whereas U2 reads averaged only 0.7% in input samples.

**Fig. 7.**
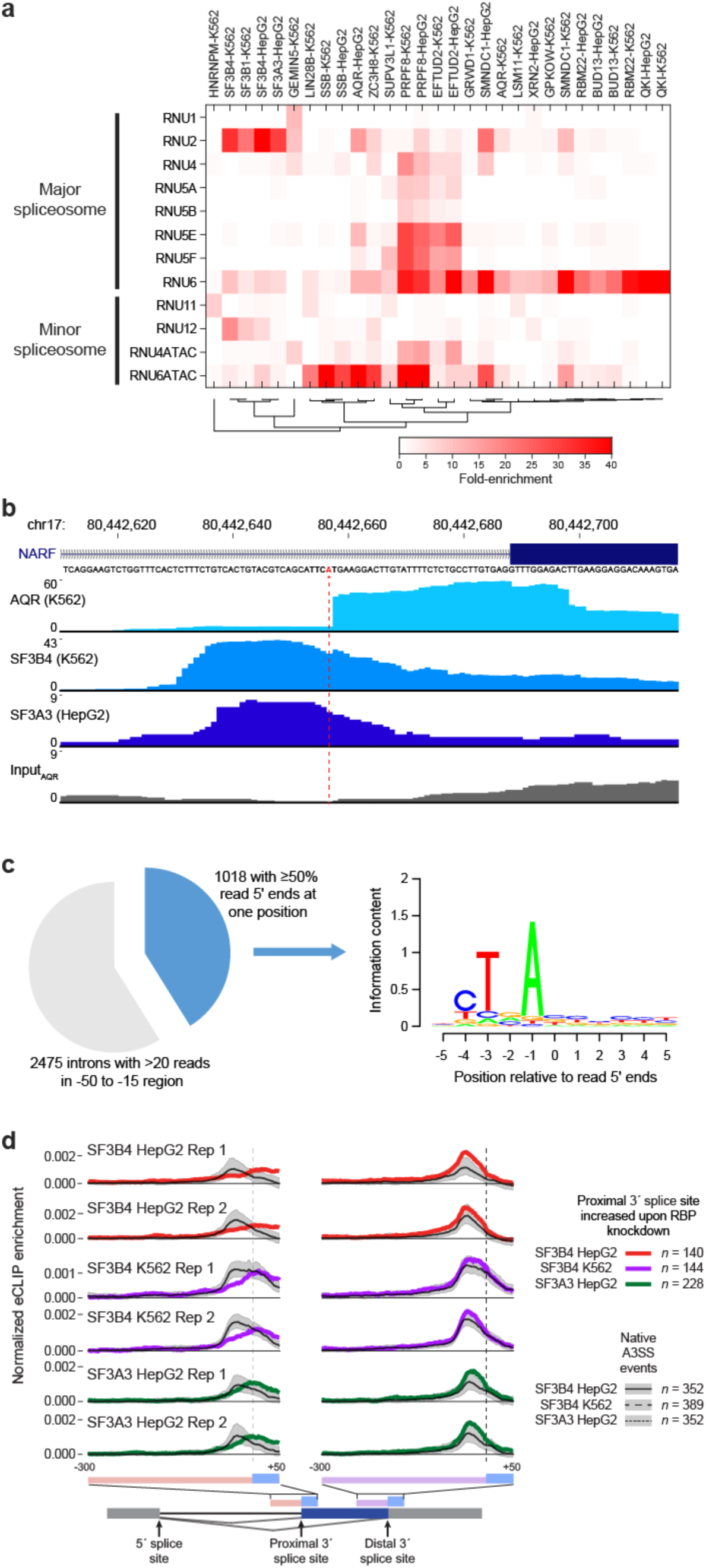
Insights from eCLIP of spliceosome-associated RBPs. (a) Heatmap indicates fold-enrichment for individual snRNAs within eCLIP datasets. Shown are all RBPs with greater than 5-fold enrichment for at least one snRNA. (b) Browser shows read density for eCLIP of AQR (K562), SF3B4 (K562), and SF3A3 (HepG2) for the *NARF* exon 11 3’ splice site region. Dotted line indicates position of enriched reverse transcription termination at crosslink sites. (c) (left) Pie chart shows all (n=2475) introns with >20 reads in the −50 to −15 (branch point) region in AQR K562 eCLIP. Blue indicates putative branch points (the subset with more than 50% of read 5’ ends at one position). (right) Motif information content for 11-mers centered on the putative branch points. Image generated with seqLogo package in R. (d) Lines indicate mean normalized eCLIP enrichment in IP versus input for SF3B4 and SF3A3 at (red/purple/green) alternative 3’ splice site extensions in RBP knockdown or (black) alternative 3’ splice site events in control HepG2 or K562 cells. The region shown extends 50 nt into exons and 100 nt into introns.

Interestingly, while many factors showed similar association between analogous snRNAs in the major and minor spliceosomes (such as PRPF8 and SMNDC1 with U6 and U6atac and SF3B1 and SF3B4 with U2 and U12), some RBPs were specifically associated with either the major (SF3A3, which was 29.5-fold enriched for U2 but 1.2-fold depleted for U12 in HepG2, and QKI, 118.6-fold enriched for U6 but 2.4-fold depleted for U6ATAC) or minor spliceosome (HNRNPM, which was 8.1-fold enriched in K562 and 7.6-fold in HepG2 for U11 but 5.3- and 4.2-fold depleted for U1) (Fig. 7a, Supplemental Fig. 7a-d). Although preliminary analysis did not show altered splicing upon HNRNPM knockdown specifically at U11/U12 introns, previous studies have suggested that HNRNPM may contribute to minor intron splicing through interactions with FUS [45].

In the first catalytic step of intron splicing, a transesterification step joins the 5’ splice site with the branch point to create an intron lariat structure (Sup. Fig. 7e). This is an essential step in splicing and helps to define 3’ splice site choice, but identification of branch points has remained challenging due to variable positioning (ranging from 20-40 nucleotides upstream of the 3’ splice site) and a degenerate sequence motif [46]. Recent efforts to use either specialized library preparation protocols or focused analysis of deep sequencing to identify branch points via lariat junction-spanning reads have enabled the identification of tens of thousands of branch points, but the regulation of branch point recognition and its role in splicing regulation remains poorly understood. Considering the RBPs profiled here, we observe multiple RBPs showing specific enrichment at branch points, including both known regulators (such as SF3 complex components SF3B4 and SF3A3), as well as novel factors (including RBM5). However, we were particularly intrigued by the observation of a striking pattern of both 5’ splice site and branch point enrichment for the RBP AQR (Fig. 7b). Knockdown of AQR yielded over 30,000 altered alternative splicing events, by far the most of any knockdown performed by the ENCODE consortium to date (including canonical splicing components including U2AF1/2 and SF3B4) [20], consistent with previous studies that indicate a role for AQR in pre-mRNA splicing [47]. However, closer inspection revealed that unlike the canonical peak shape in the branch point region observed for SF3B4 and SF3A3, the 5’ end of AQR eCLIP reads often piled up at specific positions (Fig. 7b). Using simple criteria to identify candidate branch points as positions with more than 50% of read 5’ ends within the overall −15 to −50 region, out of 2475 introns with at least 20 reads mapping to the entire branch point region we identified 1018 candidate branch points in K562 (Fig. 7c). Motif analysis of these positions yielded the canonical branch point motif signal (with 92% containing an A at the base prior to read starts) (Fig. 7d). Thus, these results suggest that AQR eCLIP signal is derived from introns after lariat formation, where reverse transcription is incapable of reading through the branch point adenosine (Sup. Fig. 7e), and that deeper sequencing of AQR eCLIP (potentially with improved methodology to enrich reads at the 3’ rather than 5’ splice site) will provide direct identification of branch points in human.

Next, we considered eCLIP signal at alternatively spliced cassette exons. Considering ‘native’ cassette exons in wild-type K562 and HepG2 cells, we observed that branch point factors SF3B4 and SF3A3 showed decreased signal at alternative exons relative to constitutive exons, consistent with U2AF2 and other spliceosomal components and potentially reflecting overall lower spliceosomal occupancy (Sup. Fig. 7f). However, at alternative 3′ splice sites with the proximal site increased upon knockdown of branch point components SF3B4 and SF3A3, we observed that average eCLIP enrichment for SF3B4 and SF3A3 was decreased at the typical branch point location but increased towards the 3’ splice site (compared to eCLIP signal at native A3SS events which utilize both distal (upstream) and proximal 3’ splice sites in control shRNA datasets) (Sup. Fig. 7g-h). Consistent with previous mini-gene studies showing that 3’ splice site scanning and recognition originates from the branch point and can be blocked if the branch point is moved too close to the 3’ splice site AG [48], these results provide further evidence that use of branch point complex association to restrict recognition by the 3’ splice site machinery may be a common regulatory mechanism [49] (Fig. 7d).

### Clustering of RBP binding identifies known and novel co-associating factors

Large-scale RBP target profiling using a consistent methodology enables cross-comparison between datasets. Considering simple overlap between peak sets for all profiled RBPs, we observed significant overlap for many pairs of RBPs, which often formed co-associating groups (Fig. 8a, left). These groups of RBPs with highly overlapping peaks generally segregated into four major categories. First, we observe high similarity between the same RBP profiled in HepG2 and K562 (including QKI, PTBP1, and LIN28B) (Fig. 8a, green). Indeed, we observe an average peak overlap of 30.0% between the same RBP in K562 and HepG2 versus 4.9% for random RBP pairings (6.1-fold increased), confirming the broad reproducibility of binding across cell types (Fig. 8b). Second, we observe many cases of high overlap between eCLIP for homologous RBPs within the same family, including TIA1 and TIAL1, IGF2BP1/2/3, and fragile X-related FMRP, FXR1, and FXR2 (Fig. 8a, yellow). Third, we observe clusters containing known co-regulating RBPs, including recognition and processing machinery for the 3’ splice site (U2AF1 and U2AF2), branch point (SF3B4 and SF3A3), and 5’ splice site (EFTUD2, RBM22, PRPF8, and others), as well as a group of RBPs that play general roles in binding the 5’UTR of nearly all genes to regulate translation (DDX3X, EIF3G, and NCBP2) (Fig. 8a, red).

**Fig. 8.**
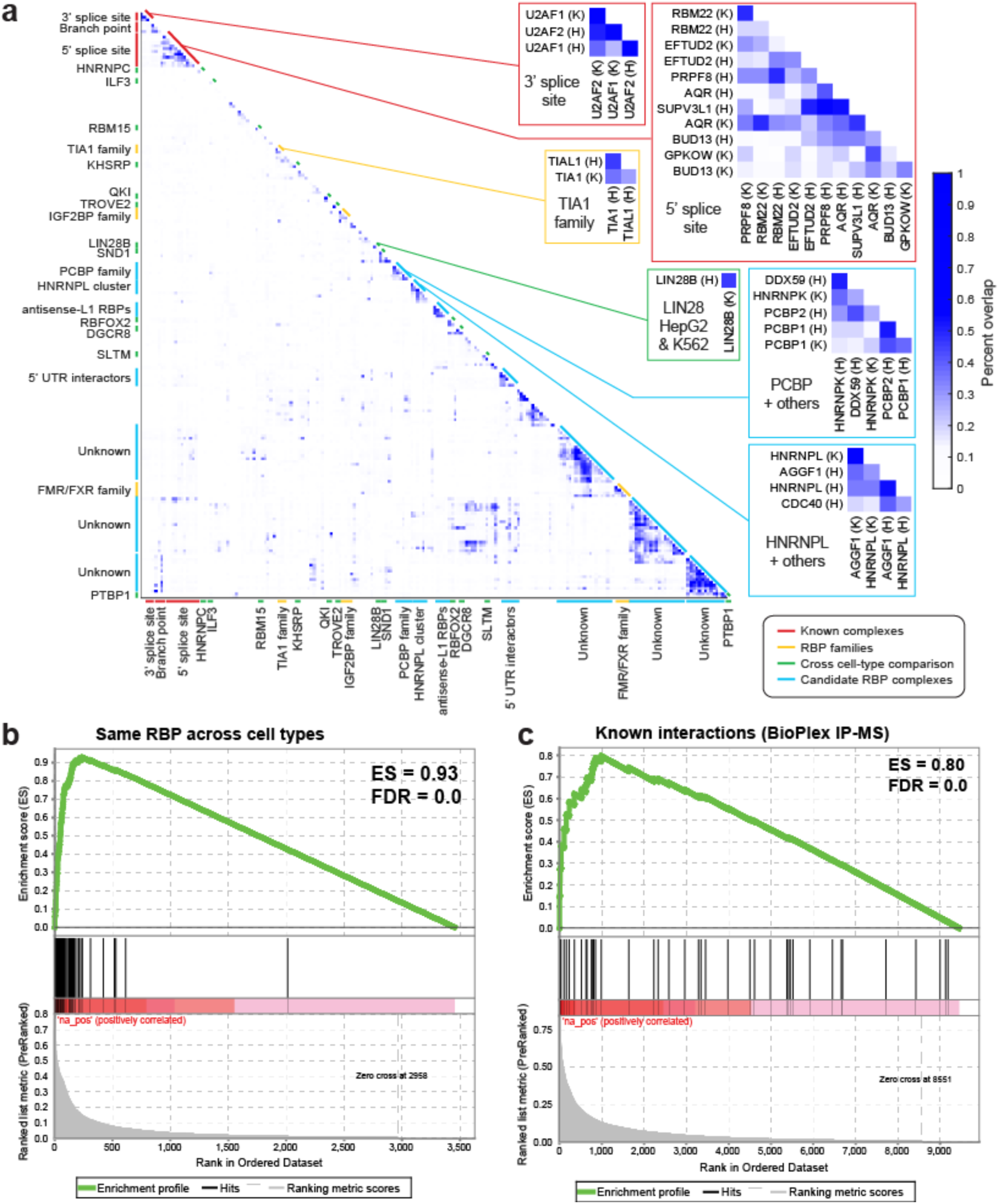
RBP co-association predicts known and novel RNP complexes. (a) Heatmap indicates the pairwise fraction of eCLIP peaks overlapping between datasets. Callout examples are shown for known complexes, RBP families, same RBP profiled across cell types, and putative novel complexes. (b) GSEA analysis comparing the fraction overlap observed profiling the same RBP in both K562 and HepG2, compared against random pairings of RBPs (with one profiled in K562 and the other in HepG2). (c) As in (b), but using the set of RBPs with interactions reported in the BioPlex IP-mass spectrometry database [50].

Interestingly, we observe unexpected clusters that suggested potential novel complexes or co-interacting partners (Fig. 8a, blue). Some clusters likely reflect overlapping targeting to specific types of RNAs: for example, one cluster contains three RBPs we described above to show specific enrichment at antisense L1/LINE elements (HNRNPM, BCCIP, and EXOSC5). The patterns of other clusters are often less clear, with some containing both well-studied RBPs as well as those with no known RNA processing roles (for example, high overlap between HNRNPL and AGGF1 across both cell types). To consider whether these likely reflected true instances of RBP co-interaction, we asked whether RBPs that had higher peak overlap were more likely to have interactions from large-scale IP-mass spectrometry experiments. Using the BioPlex 2.0 database of ∼56,000 interactions [50], we observed that RBPs with IP-MS interactions showed an average 2.3-fold increase in eCLIP peak overlap (11.4% versus 4.9% for RBPs without interactions), suggesting that there is a general correlation between peak overlap and RBP interactions (Fig. 8c).

Finally, we performed co-immunoprecipitation (co-IP) studies focusing on one predicted novel interaction group involving HNRNPL and AGGF1. We observed that AGGF1 co-immunoprecipitated HNRNPL, unlike unrelated factors RBFOX2 or FMR1 (Sup. Fig. 9a). We note that this co-IP was observed using less stringent co-IP wash buffers, but was not observed using the high salt wash buffers present in eCLIP (Sup. Fig. 8b), indicating that the overlap in eCLIP binding likely reflects independent crosslinking events to the distinct RBPs. Thus, these results indicate that the eCLIP data resource reveals many novel RBP interactions that are likely to reflect previously unidentified regulatory complexes.

## Discussion

The ENCODE RNA binding protein resource contains 1,223 replicated datasets for 356 RBPs, including *in vivo* targets by eCLIP, *in vitro* binding motifs by RNA Bind-N-Seq, subcellular localization by immunofluorescence, factor-responsive expression and splicing changes by knockdown/RNA-seq, and DNA associations by ChIP-seq [20]. This unique resource has already proven useful in characterizing allele-specific RBP interactions [51, 52], identify candidate regulators of miRNA processing [53], predicting whether RNAs are protein-coding or non-coding [54], and identifying novel factors which act to suppress improper RNA processing caused by retrotransposable elements [36], and will continue to enable researchers to ask broad questions about basic RNA processing mechanisms, deeply consider the functional roles of an individual RBP, or even query an RNA of interest in order to gain insight into potential regulators. Here we describe examples how integrated analyses of binding profiles obtained from eCLIP can yield novel insights into both processing of standard mRNAs as well as other RNA families, including identifying new characteristics of ribosomal RNA processing and the role of RBP interactions with retrotransposable elements.

### Inference of RBP function based on eCLIP enrichment patterns

Deep profiling of RBPs associated with a specific RNA processing pathway can yield unique insights into the specialization of RBPs. For example, profiling of 30 RBPs associated with RNA degradation gave insights into specific RNP complex variants with roles targeting specific subtypes of RNAs, providing a comprehensive view of how the wide array of RNAs in the cell are turned over [8]. In contrast, the relatively unbiased selection of 150 RBPs profiled here enabled us to query across a wide variety of RBP functions and binding modalities and, at a broad level, address the basic question of whether RNA targets identified by CLIP can generally predict the likely function of the RBP of interest. This analysis confirmed that both the RNA transcript class level, where eCLIP enrichment for ribosomal RNA or retrotransposable elements correlated with specific RBP functions focused around these element types, as well as at the regulatory region level, where enrichment at 5’UTRs or branch point regions corresponded to specific RBP functional roles.

Although these overall patterns match well with our existing understanding of RBP functions, the validation of distinctive profiles for different functions enables deeper interpretation of RBPs based solely on eCLIP. For example, we observed specific enrichment for GEMIN5 beginning in the 5’UTR and peaking at the start codon, providing further genome-wide validation for the role of GEMIN5 in translation regulation [55] (Fig. 2B). Similarly, the association of ZC3H11A at the 3’ end is consistent with iCLIP signal observed for TREX complex component ALYREF [56] and provides further transcriptome-wide evidence to support the observation that ZC3H11A plays an essential role in export of polyadenylated mRNA through interaction with the TREX complex [57]. As we continue to profile additional RBPs, these results suggest it should become possible to predict RBP function with increasing resolution based on association patterns.

Considering meta-exon plots focused on exon/intron boundaries, we observe expected patterns of eCLIP enrichment at canonical splicing elements (5’ and 3’ splice sites and branch points). We also observe classes of RBPs with broader patterns of enrichment, with a particularly interesting group showing a stereotypical pattern of high enrichment at the 5’ end of introns (extending hundreds of bases into the intron). Notably, this cluster contains multiple factors with links to co-transcriptional RNA processing, including CSTF2T [58], XRN2 [59] and Nono [60], suggesting that this group may reflect interactions that mark the time period between 5’ splice site transcription and splicing. Interestingly, this cluster also contains FET family proteins FUS and EWSR1, consistent with previous CLIP-seq studies which identified a similar ‘sawtooth’ pattern for FUS [61] and suggesting that co-transcriptional deposition may be a general regulatory principle for this family of neurodegenerative disease-associated RBPs.

### Enrichment patterns reveal insights into ribosomal RNA processing

The enrichment for previously identified rRNA processing factors suggests that many additional factors here may represent unexplored regulators. Indeed, building upon rRNA enrichments observed from the analyses described here, further research has led to validation of NOL12 [62] and AATF [63] as novel regulators of ribosomal RNA processing, indicating that there remain more RBPs with unexplored roles in ribosomal RNA processing.

Another benefit of the unbiased approach presented here is that it enables identification of novel potential sites of regulatory activity, as our analysis of the 45S ribosomal RNA precursor indicates a surprising cluster of substantial RBP eCLIP enrichment at an uncharacterized region located between the A0 and 1 processing sites. This region (particularly the sharp peak observed in LIN28B eCLIP) is centered on a putative ribosomal-encoded microRNA (rmiR-663) [31], and our analysis indicates that the reads do appear to be derived from ribosomal RNA rather than paralogous genomic-encoded microRNAs. However, we do not observe enrichment in DROSHA or DGCR8 eCLIP in this region (Fig. 3e), suggesting that rmiR-663 does not progress through the normal miRNA maturation pathway. Thus, it remains unclear whether this represents a bona fide microRNA, or more complex regulation of either ribosomal RNA processing or maturation of other microRNAs. Indeed, LIN28B has previously been shown to inhibit let-7 biogenesis by sequestering primary let-7 transcripts in the nucleolus away from DROSHA processing [30]. Although one model could be that LIN28B association to this region simply is an artifact of nucleolar localization, the high abundance of 45S rRNA overall (and nearly 500-fold enrichment for LIN28B at this site) suggests that the rmiR-663 region might instead act to sequester LIN28B, thereby coupling LIN28B inhibition of let-7 microRNA biogenesis to ribosomal RNA transcription and abundance. Similarly, although SSB has previously been associated with microRNA processing through interactions with pre- and pri-miRNAs [64], SSB traditionally interacts with RNA Polymerase III transcripts [65], potentially suggesting distinct Polymerase III transcription of this region in addition to Polymerase I transcription of the entire 45S transcript. Further work will be required to fully confirm whether rmiR-663 is actually processed from the 45S to maturity as a functional miRNA incorporated into the RISC complex for mRNA targeting, or whether these other potential regulatory modalities act to control other aspects of rRNA or microRNA processing.

### Retrotransposable element suppression: a major function for many RBPs

Analysis of Alu elements identified 3 RBPs with at least 4-fold enrichment, each of which appears to reflect a different underlying mechanism. The most enriched RBP, HNRNPC, has previously been shown to suppress cryptic 3’ splice site signals in antisense Alu elements [35]. In contrast, ILF3 (enriched for both sense and antisense Alu elements) has previously been shown to interact with RNA editing mediator ADAR1 [66], and the majority of ADAR1 targets and edited sites throughout the genome occur at Alu elements [67]. Further research has now revealed that ILF3 knockdown induces RNA editing, and suggested that ILF3 binding to Alu elements generally acts to repress RNA editing at these sites [37]. The third RBP, RNA Polymerase II subunit POLR2G, may reflect previous observations of antisense L1 and (particularly inverted tandem) Alu elements repressing PolII progression [38, 68]. Indeed, we observe that POLR2G eCLIP shows enrichment for sense Alu (2.3-fold), sense L1 (1.8-fold), and antisense L1 (4.0-fold) elements as well as antisense Alu (5.0-fold), providing further evidence that the high propensity for such regions to form structural elements may generally inhibit polymerase progression through these regions, leading to increased dwell time for POLR2G.

Similarly, analysis of L1 element enrichment revealed multiple modalities of regulatory activity. One function of RBP association to L1 is to suppress retrotransposition activity, and indeed we observed that three RBPs (PPIL4, SAFB, and TRA2A) showed both eCLIP enrichment for sense L1 elements and act to suppress L1 retrotransposition activity in genome-wide screening data. For RBPs enriched for antisense L1 elements, we instead see signatures of RBPs acting to increase RNA expression, extending a similar analysis recently published (that included an earlier release of the ENCODE eCLIP resource along with other iCLIP datasets) that revealed widespread association with L1 elements by RBPs [36]. From these and other works, it is now becoming clear that suppression of aberrant RNA processing due to retrotransposable elements is a major responsibility of many RNA binding proteins, suggesting that the genome has evolved to devote substantial resources to this effort.

### Large-scale RBP target maps provide unique opportunities for further specialized insights

It is notable that the above enriched RNA element classes often reflected a substantial fraction of eCLIP reads, suggesting that they may represent dominant functions of the RBP. For example, antisense L1 elements constituted 19-27% of eCLIP reads for HNRNPM and MATR3 and antisense Alu elements were 13-18% of reads in HNRNPC eCLIP. Similarly, 42-56% of UTP18, 27-31% of WDR43, and 16% of HepG2 LIN28B eCLIP reads mapped to the 45S ribosomal RNA precursor. Thus, these results strongly argue that analysis of CLIP data should include proper quantitative analysis of reads mapping to non-mRNA regions, as they can in some cases represent the dominant binding modality of the RBP and should be considered in interpreting potential functional roles of the RBP in regulating RNA processing.

Intriguingly, we even observed significant differences even between RBP components of the same RNP complex. For example, 41.0% of SF3A3 HepG2 eCLIP reads mapped to RNU2 snRNA versus only 8.5% mapping to proximal intronic regions; in contrast, SF3B4 was far more even (23.1% proximal intronic in HepG2 and 17.8% in K562, versus 17.0% and 19.7% RNU2 in HepG2 and K562 respectively). Although we cannot rule out that this difference in crosslinking to snRNA versus intron reflects underlying amino acid biases in UV crosslinking efficiency, it does confirm that CLIP profiling of multiple RBP members of an RNP complex can yield distinct insights into interaction patterns and regulatory roles of the complex, suggesting that it is critical to assay multiple independent proteins to gain a full understanding of the target repertoire of an RNP complex.

In addition to specific insights into the RBPs themselves, we anticipate that the broad diversity of RBPs profiled and RNA elements and features bound will spur further development of methods targeted towards specific RNA processing steps. For example, the peak distribution pattern of the CDS.5P class (and RPS3 in particular) resembles the average profile observed using ribosome profiling [69], suggesting that RPS3 eCLIP may capture ribosome association on translating mRNAs and could be used as a general approach to assay translation. Similarly, our meta-exon analysis of AQR (followed by further analysis of crosslink-induced termination sites) showed that AQR eCLIP could identify branch points for a set of highly abundant introns, suggesting that further development of profiling of AQR binding targeted to 3’ splice site regions could yield a highly specific approach to identification of branch points transcriptome-wide.

The diversity of distinct RBP association patterns can also be flipped to predict features of a queried RNA. For example, recent work used the ENCODE eCLIP resource to identify UPF1 as one of many RBPs with specific enrichment at 3’UTRs [54]. This finding enabled improved prediction of whether a queried transcript was a protein coding versus long non-coding RNA by incorporating presence (or absence) of UPF1 eCLIP signal as a biomarker for translation [54]. Similarly, our unbiased analysis of foci of enrichment on the 45S rRNA precursor suggested two regions as notably highly enriched across multiple RBPs, one of which matches a well-characterized region (between the canonical 01 and A0 processing sites) with another suggesting interesting regulatory mechanisms linking ribosomal RNA and microRNA processing. Similar analysis identifying eCLIP datasets with enrichment on regulatory noncoding RNAs *Xist* and *Malat1* also suggested that the patterns of RBP enrichment often correlate with specific structural and functional domains on these noncoding RNAs [18]. With the continuing release of profiles for additional RBPs, we expect that identification of these distinct RBP ‘states’ may serve as a useful method for independent prediction of key regulatory domains within these non-coding RNAs.

## METHODS

### eCLIP datasets used

Enhanced CLIP (eCLIP) datasets used were obtained from the ENCODE data coordination center (https://www.encodeproject.org) with accession identifiers listed in Supplementary Table 1. Unless otherwise indicated, standard peak analysis used the set of peaks identified as irreproducible discovery rate (IDR) reproducible and meeting fold-enrichment (≥8-fold) and significance (*p-*value ≤ 10^-3^) in immunoprecipitation versus paired size-matched input. RNA binding protein function annotations and localizations were obtained from [20] (Supplementary Table 2). The list of RNA binding proteins was obtained from [3]. The list of transcription factors was obtained from [70], using the “a”, “b”, and “other” classes.

### Biotin-based visualization of RBP-coupled RNA

A step-by-step version of the biotin-based labeling protocol is available at https://www.protocols.io/view/biotin-labelling-of-immunoprecipitated-rna-v1pre-7z4hp8w. In brief, for visualization experiments, HepG2 or K562 cells were prepared identically to eCLIP experiments up until the first RNA adapter ligation: 20 million cells were lysed in 1 mL 4°C eCLIP lysis buffer, fragmented for 5 min at 37°C with 40U RNase I (Ambion), centrifuged at 15k RPM for 3 minutes at 4°C (with supernatant kept) to clear lysate, and incubated with rotation overnight with antibody coupled to species-specific secondary beads (10 μg primary antibody as indicated coupled to 125 uL of Sheep anti-Rabbit or anti-Mouse Dynabeads; ThermoFisher). After incubation, samples were washed once with eCLIP wash buffer, washed twice with high salt wash buffer, and washed three times with wash buffer. FastAP and T4PNK reactions were performed on-bead as previously described for eCLIP, followed by one wash with high salt wash buffer and 3 washes with wash buffer. At this point, a modified RNA linker ligation was performed with standard eCLIP ligation conditions (buffer and High Concentration T4 RNA Ligase) but with 500 pmol pCp-Biotin (Jena Bioscience) in place of the RNA adapter, and samples were incubated at 16°C. For some experiments, immunoprecipitations were performed on 4 million cells; for these experiments, half reactions were used for the pCp-biotin ligation step. After ligation, samples were washed once with high salt wash buffer and three times with wash buffer, followed by standard SDS-PAGE electrophoresis and transfer to nitrocellulose membranes. Visualization was performed using the Chemiluminescent Nucleic Acid Detection Module Kit (ThermoFisher), following manufacturer instructions for blocking, washes, and labeling. Imaging was performed on the Azure C600 platform. For ^32^P experiments, radiolabeling was performed as previously described [71].

### Family-aware mapping to multi-copy elements

The software pipeline used to quantify enrichment for retrotransposable and other multi-copy elements is available at https://github.com/YeoLab/repetitive-element-mapping, and was initially described in [20] but is described in more complete detail below. This release includes scripts, detailed documentation, and database files necessary to perform the described analyses.

A database of multicopy elements was generated based on 5606 transcripts obtained from GENCODE v19 covering 34 families of abundant non-coding, multicopy, and other types of RNA refractory to standard peak analysis, including families within the broader rRNA (RNA18S, RNA28S, RNA5S, RNA5-8S), snoRNA (SNORD, SNORA, RNU105, RNU3, RNU7, snoU13, snoU109, U8), snRNA (RNU1, RNU2, RNU4, RNU4ATAC, RNU5A, RNU5B, RNU5D, RNU5E, RNU5F, RNU6, RNU6ATAC, RNU11, RNU12), vault RNA (VTRNA1, VTRNA2, VTRNA3), non-coding RNA (H1RNA, RN7SK, RN7SL, MRP, YRNA), and small Cajal body-specific RNA (SCARNA) broader classes (Supplemental Table 3). Each family contained GENCODE v19 annotated transcripts as well as their pseudogenes. To this set were added a family for tRNAs (606 tRNA transcripts were obtained from GtRNAdb [72], and each tRNA was included in two versions: one variant including 50nt of genome flanking sequences, and one mature variant that included the canonical CCA tail), mitochondrial transcripts (which were initially added as one class of 37 annotated genes, but ultimately counted as two families based on H- or L-strand position that included not only gene-mapping reads, but also intergenic reads mapping uniquely to the mitochondrial genome), the rRNA RNA45S precursor transcript (NR_046235.1, obtained from GenBank), a ‘simple repeat’ class containing 501 60-mer sequences containing simple repeats of all 1 to 6-nt k-mers, and 49 families comprising 705 total human repetitive elements obtained from the RepBase database (v. 18.05)[73]. Within each family, transcripts were given a priority value, with primary transcripts prioritized over pseudogenes. Mapping to the reverse strand of a transcript was counted separately from forward strand mapping, creating a second ‘antisense’ family for each RNA family above (which utilized the same element priority order), with the exception of simple repeats (which were all combined into one family).

To quantify eCLIP signal, paired end sequencing reads were first adapter trimmed as previously described [18]. Next, reads were mapped against the repetitive element database using bowtie2 (v. 2.2.6) with options “-q --sensitive -a -p 3 --no-mixed –reorder” to output all mappings. Read mappings were then processed as follows. First, for each paired-end read pair only mappings with the lowest alignment scores summing both mismatch penalties (defined as *MN + floor((MX-MN)(MIN(Q, 40.0)/40.0))* where Q is the Phred quality value, and default values MX = 6, MN = 2, as described in bowtie2 reference material) and gap penalties (defined as *GO + N * GE*, where GO = gap open = 5, GE = gap extend = 3, N = gap length) were kept. Next, the mapping to the transcript with the highest priority within a RNA family (as listed above) was identified as the ‘primary’ match mapping. At this stage, read pairs which had equal best alignments to multiple repeat families were discarded, with only reads mapping to a single repeat family considered for further quantification.

Next, these RNA family mappings were integrated with unique genomic mapping from the standard eCLIP processing pipeline (using read mapping prior to PCR duplicate removal). For read pairs that mapped both to an RNA family above as well as uniquely to the genome, the mapping scores (as defined above) were compared. If the unique genome mapping was more than 2 mismatches per read (24 alignment score for the read pair) better than to the repeat element, the unique genomic mapping was used; otherwise, it was discarded and only the repeat mapping was kept. Next, PCR duplicates were removed by comparing all read pairs based on their mapping start and stop position (either within the genome or within the mapped primary repeat) and unique molecular identifier sequence, and all but one read pair for read pairs sharing these three values were defined as PCR duplicates and removed. At this stage, RepeatMasker-predicted repetitive elements in the hg19 genome were additionally obtained from the UCSC Genome Browser [23]. Element counts for RepBase elements were therefore determined as the sum of repeat family-mapped read pairs (described above) plus the number of reads that mapped uniquely to the genome at positions which overlapped (by at least one base) RepeatMasked RepBase elements. Reads uniquely mapping to non-RepBase genomic regions were then annotated into one of 11 additional classes in the following priority order (based on GENCODE v19 annotations): CDS, 5′UTR and 3′UTR, 3’UTR, 5′UTR, proximal intronic (within 500nt of splice sites), distal intronic (remaining intronic regions), non-coding exonic, non-coding proximal intronic, non-coding distal intronic, antisense to gencode transcripts, and intergenic.

Finally, the number of post PCR-duplicate removal read pairs mapping to each class was counted in both IP and paired input sample and normalized for sequencing depth (using the total number of post-PCR duplicate read pairs from both unique genomic mapping as well as repeat mapping as the denominator to calculate fraction of reads). Significance was determined by Fisher’s Exact test, or Pearson’s Chi-Square test if all expected and observed values were five or more. Relative information content of each element in each replicate was calculated as 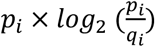, where *p_i_* and *q_i_* are the fraction of total reads in IP and input respectively that map to element *i*. To combine two biological replicates, the average reads per million (RPM) was calculated across two IP samples and compared against the paired input experiment to calculate one overall fold-enrichment and relative information value per dataset.

### Validation of RNA element links with RBP functional annotations

To quantify whether RNA element enrichment matched with RBP functions, a set of positive control pairings were generated between RNA elements with known links to either RBP function or known RBPs contained within a well-characterized ribonucleoprotein complex (Sup. Fig. 2a). 140 datasets for which the RBP had at least one of these annotated functions were chosen, and were considered as ‘true positives’ if the most-enriched RNA element class (by relative information) matched this known RBP function. Datasets were sorted by relative information of the most-enriched class. Accuracy (defined as *(TP + TN) / (TP + TN + FP + FN)*) was then calculated for each sequence relative information cutoff, and the maximum point (0.2) was chosen.

### Ribosomal RNA analysis

RBPs with roles in ribosomal RNA processing were obtained from [27]. Position-wise relative information was calculated as above, using the number of reads overlapping the position in IP versus input for each dataset. To obtain a cutoff for further analysis, RBPs were sorted by the maximum position-wise relative information on the 45S rRNA precursor, and at each value the F1 score was calculated (defined as *(2 × TP) / (2 × TP + FP + FN)*). The maximum point at 0.064 was used for further analysis.

To quantify enrichment at the rmiR-663 ribosomal versus genomic paralog loci, sequences of rmiR-663 and four genomic-encoded paralogs (miR-663a, miR-663b, AC010970.1, and AC136932.1) were obtained from the UCSC genome browser, along with 100nt of flanking sequence. Only reads that perfectly aligned (with zero mismatches or gaps) to these sequences were counted for further analysis.

### Retrotransposable element analysis

L1 retrotransposition genome-wide CRISPR screening data was obtained from Liu *et al*. [39], using Combo casTLE Effect scores from K562 cells. Bonferroni correction was performed on uncorrected casTLE p-values using n=15 (the number of L1 (sense)-enriched RBPs queried).

To calculate change in expression of L1-containing bound genes, DESeq-calculated gene expression fold-changes for RBP knockdown/RNA-seq data were obtained from the ENCODE DCC (http://www.encodeproject.org) for all RBPs with both eCLIP and RNA-seq performed in the same cell type. L1 sense and anti-sense elements were taken from RepeatMasker-predicted repetitive elements in the hg19 genome obtained from the UCSC Genome Browser [23]. For each gene in Gencode v19, the transcript with the highest abundance in rRNA-depleted total RNA-seq in HepG2 (ENCODE accession ENCFF533XPJ, ENCFF321JIT) and K562 (ENCFF286GLL, ENCFF986DBN) was chosen as the representative transcript, and the set of expressed genes (10,247 in HepG2 and 9,162 in K562 with TPM ≥ 1) were considered. Next, genes were separated into three classes: ‘≥ 1 bound L1(as)’ genes with at least one antisense L1 element that overlapped a significant peak identified in eCLIP, ‘bgd with ≥ 1 L1(as)’ genes with at least 1 antisense L1 element but did not have an element that overlapped with an eCLIP peak, or ‘Bgd’ which contained all expressed genes. Significance was determined by Kolmogorov-Smirnov test with no multiple hypothesis testing correction.

To compare reference versus divergent L1 elements, we defined ‘canonical’ reads as those which mapped best (and were assigned) to sequences present in RepBase, whereas ‘divergent’ reads mapped better to unique genomic loci than to the reference sequence.

Calculation of overall element coverage (Sup. Fig. 4b) was based on the above set of 9,162 reference transcripts in K562 expressed with TPM ≥ 1.

### Meta-gene and meta-exon peak density maps

To generate meta-gene and meta-exon maps, for each gene in Gencode v19, the transcript with the highest abundance in rRNA-depleted total RNA-seq in HepG2 (ENCODE accession ENCFF533XPJ, ENCFF321JIT) and K562 (ENCFF286GLL, ENCFF986DBN) was chosen as the representative transcript, and the set of expressed genes (10,247 in HepG2 and 9,162 in K562 with TPM ≥ 1) were considered. Datasets with fewer than 100 mRNA-overlapping peaks were discarded, leaving 205 datasets. Next, each gene was split into 162 bins (13 for 5′UTR, 100 for CDS, 49 for 3′UTR), based on the median 5′UTR, CDS, and 3′UTR lengths of highly expressed (TPM ≥ 10) Gencode v19 transcripts in K562 cells. For each eCLIP dataset, the average peak coverage for each bin was calculated for each gene, and then averaged over all genes to generate final meta-gene plot. To generate confidence intervals, bootstrapping was performed by randomly selecting (with replacement) the same number of transcripts and calculating the average position-level peak coverage as above, with the 5^th^ and 95^th^ percentiles (out of 100 permutations) shown. For further visualization and analysis, only 104 RBPs where the 5^th^ percentile was at least 0.002 peaks per gene (∼20 peaks in at least one bin) were considered. Normalized coverage was then calculated by setting the maximum position to one and minimum position to zero for each eCLIP dataset. Cross-position correlations were calculated using normalized coverage for across all 104 RBPs at each position. Odds ratios and significance (determined by Fisher’s Exact Test, or Yates’ Chi Square test if observed and expected values were greater than five) utilized RBP annotations (Supplementary Table 2) from [20].

To generate meta-exon plots for each eCLIP dataset, for all internal exons (excluding the first and last exons) the region from 500nt upstream to 500nt downstream (for introns less than 1000nt, the region was split with half assigned to the upstream exon and half to the downstream exon) was queried for the presence of significant (IDR) peaks. Finally, the number of peaks at each position was averaged over all events to obtain the final meta-exon value. To generate confidence intervals, bootstrapping was performed by randomly selecting (with replacement) the same number of transcripts and calculating the average position-level peak coverage as above, with the 5^th^ and 95^th^ percentiles (out of 100 permutations) shown. For further analysis, only datasets with at least 100 IDR peaks were considered. Next, after calculating meta-exon profiles and confidence intervals as above, datasets that did not have at least one position with the 5^th^ percentile bootstrap value above a minimal cutoff of 0.0005 (∼5 peaks observed at that position) were discarded to leave 133 datasets for further consideration. Finally, for visualization of comparison across RBPs (Figure 6), an additional normalization was performed by dividing each position by the maximum meta-exon value for that dataset, in order to scale the meta-exon profiles between 0 and 1.

### Analysis of AQR enrichment at branch points

To identify points of enriched read termination in AQR eCLIP, regions from −50nt to −15nt from annotated 3′ splice sites were obtained from GENCODE v19, and the subset of regions with at least 20 overlapping reads in AQR eCLIP in K562 cells were taken for further analysis. Points of enrichment were identified as those where more than half of reads overlapping the overall region terminated at the same position. Motif analysis was performed by counting the frequency of 11-mers centered on the read start position with 5nt flanking on either side. Motif logos were generated with seqLogo (R).

### Enrichment of branch point factors at alternative 3′ splice site events

Splicing maps profiling normalized enrichment for SF3B4 and SF3A3 at RBP knockdown-responsive alternative 3’ splice site events were generated as previously described [20, 74]. In brief, the set of differential 3’ splice site events for RBP-knockdown/RNA-seq was identified from rMATS analysis between RBP knockdown and paired non-target control. Normalized read density in eCLIP was then calculated for each differential event by subtracting input read density from IP read density (each normalized per million mapped reads). To weigh each event equally, position-wise subtracted read density was then normalized to sum to one across the entire event region (composed of 50nt of exonic and 300nt of flanking intron), including a pseudocount of one read (normalized by total mapped read density) at each position. The highest 2.5% and lowest 2.5% values at each position across all events were then removed, and the mean was then calculated across all other events to define the final splicing map. As a control, a set of ‘native’ alternative 3’ splice site events was defined as those which showed alternative usage (0.05 < inclusion < 0.95) in control K562 or HepG2 cells respectively. Confidence intervals were generated by randomly sampling the number of events in the RBP-responsive class from the native alternative 3’ splice site set 1000 times, processing this sampled set as described above, and plotting the 0.5^th^ to 99.5^th^ percentiles.

### Co-occurrence of RBP eCLIP peaks and validation of sub-complexes of RBPs

Overlap between eCLIP datasets A and B was determined by calculating the fraction of significant and reproducible peaks in dataset A that overlapped (by at least one base) a peak in dataset B, and vice versa the fraction of peaks in B that overlapped a peak in A, and taking the maximum of those fractions as the overall pairwise fraction overlap. Only datasets with at least 100 reproducible and significant peaks were used for this analysis. Gene Set Enrichment Analysis was performed using the GSEA software package [75]. RBP interaction data was obtained from the BioPlex 2.0 dataset [50].

IP-western validation was performed using HNNRPL (ab6106, Abcam), RBFOX2 (A300-864A, Bethyl), FMR1 (RN016P, Bethyl), AGGF1 (A303-634A, Bethyl), and TNRC6A (RN033P, MBLI) antibodies in UV crosslinked K562 cells. Immunoprecipitation in high-salt wash conditions was performed using standard eCLIP wash buffers, beads, and other reagents [18]. Low-salt co-immunoprecipitation conditions used identical conditions, except for lysis buffer (50 mM Tris-HCl pH 7.5, 150 mM NaCl, 1% Triton X-100, 0.1% Sodium deoxycholate, and Protease Inhibitor cocktail (Promega)) and wash buffer (5 washes total in TBS + 0.05% NP-40). Westerns were probed with HNNRPL (ab6106, Abcam) primary antibody and TrueBlot secondary (Rockland).

## Figure legends

**Supplemental Fig. 1.**
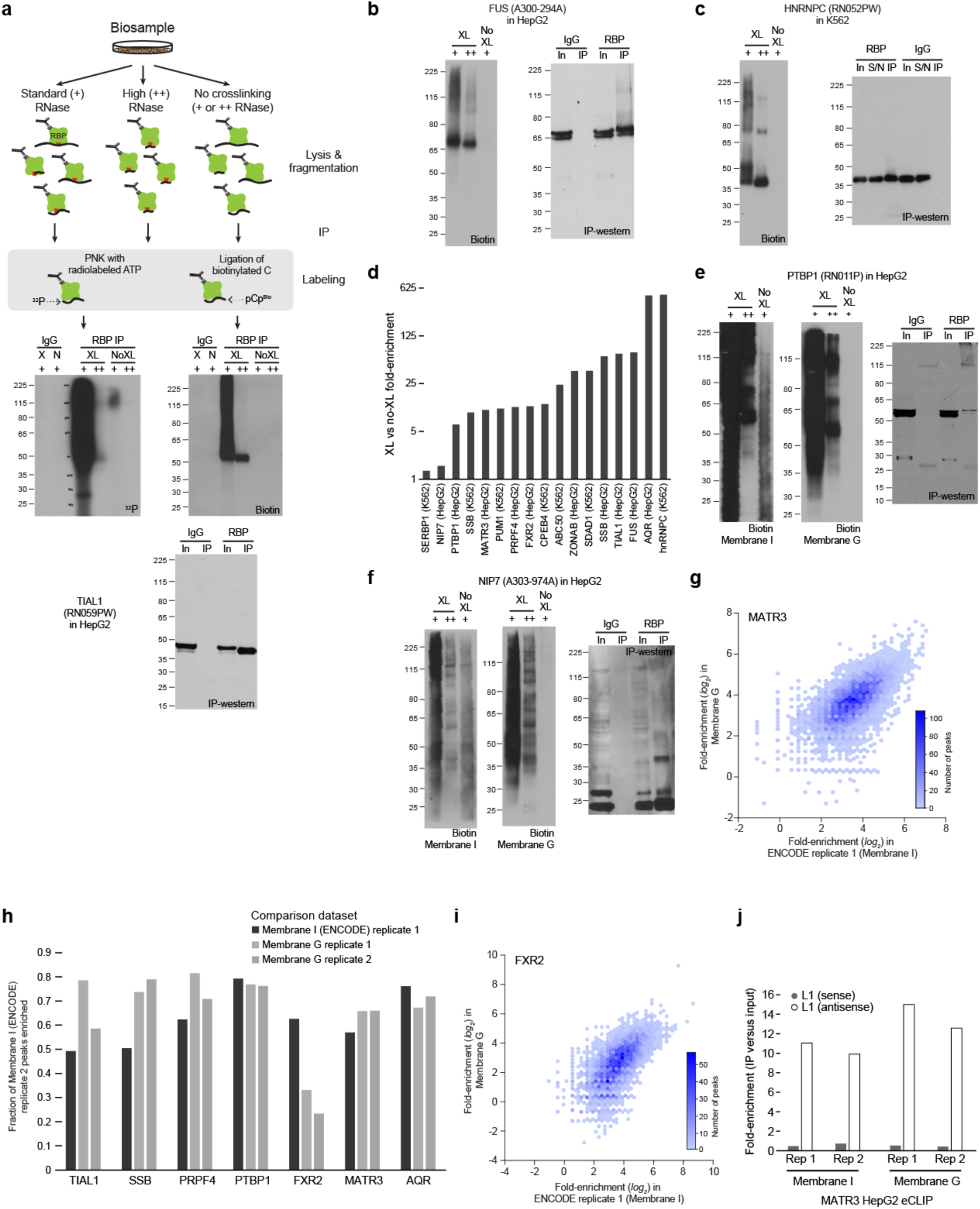
Visualization of RBP:RNA complexes with biotin-labeling. (a) (top) Schematic of RBP:RNA visualization experiments, in which three samples are subjected to immunoprecipitation: crosslinked cells with standard (40U) RNase, crosslinked cells with high (333U) RNase, and non-crosslinked cells with either standard or high RNase. RNA was then labeled either through radiolabeling with T4 PNK and [γ-32P]-ATP followed by autoradiography, or with T4 RNA Ligase and pCp-Biotin followed by chemiluminescent imaging with streptavidin-conjugated horseradish peroxidase. (bottom) Example RNA imaging with ^32^P and biotin-labeling after TIAL1 immunoprecipitation, with standard IP-western shown below. (b-c) Biotin-based RNA labeling for (b) FUS in HepG2 and (c) HNRNPC in K562. (d) Bars indicate the fold-enrichment in biotin-labeled RNA signal between crosslinked versus non-crosslinked samples (with 40U RNase) for the size range from which RNA is isolated (from protein to 75 kDa above). Shown is data from membrane I. Quantification was performed in ImageJ. (e-f) Biotin-based RNA labeling for (e) PTBP1 in HepG2 and (f) NIP7 in HepG2. For each, immunoprecipitated sample was labeled with pCp-Biotin and split in half, with one half transferred to nitrocellulose membrane from supplier I, and the other half transferred to nitrocellulose membrane from supplier G. (right) IP-western experiment from the paired eCLIP experiments). (g) Density plot indicates the number of eCLIP peaks for MATR3 in HepG2 identified as significant in ENCODE replicate 2 that have the indicated fold-enrichment in (x-axis) ENCODE replicate 1 (performed with membrane I) versus (y-axis) a new eCLIP replicate performed with membrane G. Color indicates the number of points within each hexagon. (h) Bars indicate the fraction of significantly-enriched peaks in ENCODE replicate 2 (performed with membrane I) that are also significantly enriched in (black) ENCODE replicate 1 (membrane I) or (gray) replicates 1 or 2 of a new eCLIP experiment performed with membrane G in the same cell type with the same antibody. (i) Density plot indicates the number of eCLIP peaks for FXR2 in HepG2 identified as significant in ENCODE replicate 2 that have the indicated fold-enrichment in (x-axis) ENCODE replicate 1 (performed with membrane I) versus (y-axis) a new eCLIP replicate performed with membrane G. Color indicates the number of points within each hexagon. (j) Bars indicate the fold-enrichment for read density at (white) sense or (gray) antisense L1 elements in MATR3 eCLIP in HepG2.

**Supplemental Fig. 2.**
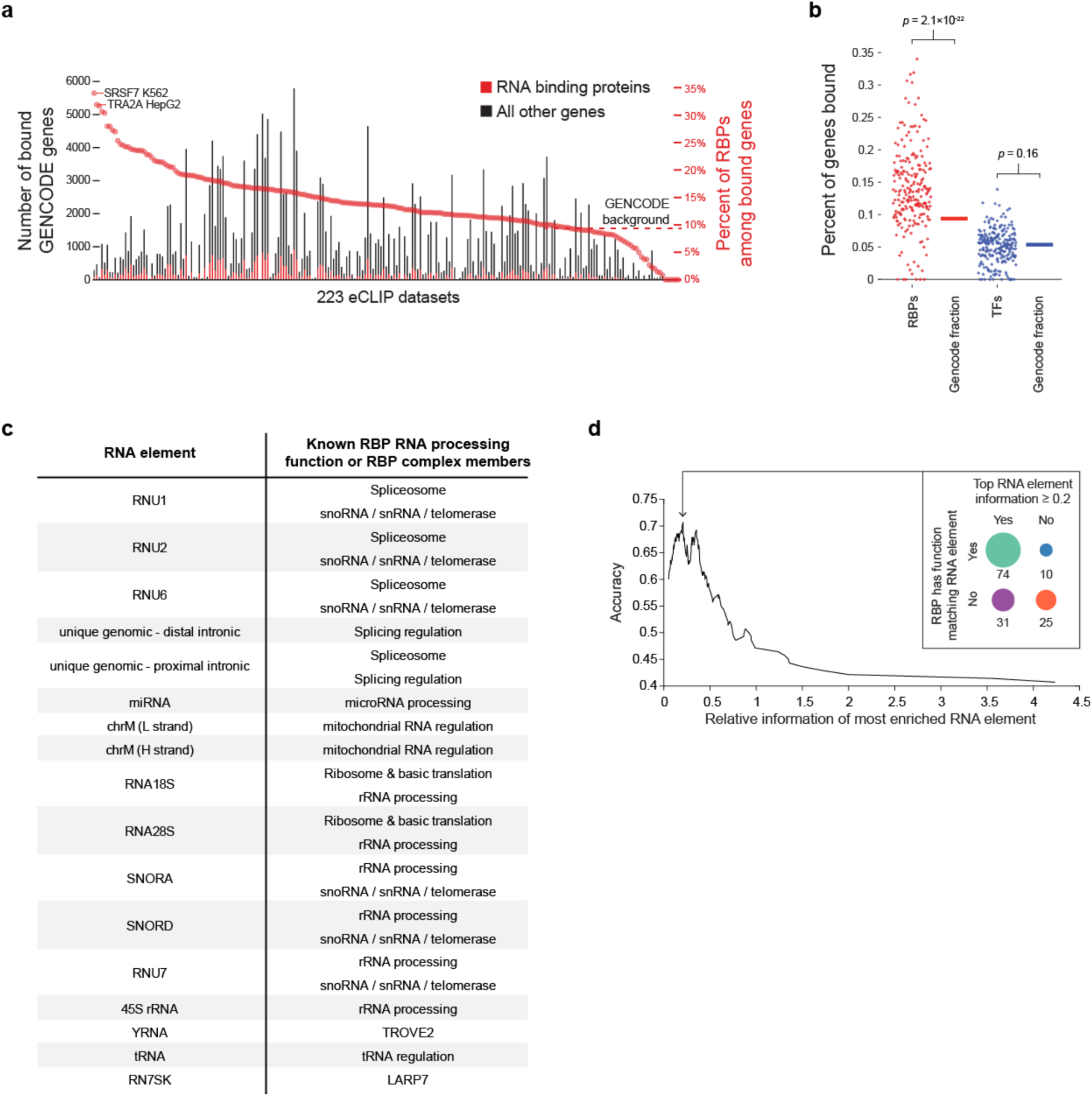
Quantification of genomic signal versus repetitive elements and other non-uniquely mapped reads. (a) Stacked bars indicate number of (red) RBPs and (black) all other GENCODE genes with eCLIP peaks in each dataset. Red circles indicate the percent of bound genes that are RBPs. eCLIP datasets are sorted by percent of RBPs. (b) Percent of genes that are (red) RBPs or (blue) transcription factors (TFs) are indicated for all 223 eCLIP datasets, compared with the fraction of RBPs out of all GENCODE genes that contain at least one peak in any of the 223 eCLIP datasets. Significance was determined by two-sided Kolmogorov-Smirnov test. (c) Table indicates RNA elements paired with canonical functional annotations for their respective ribonucleoprotein complexes, or (for YRNA and RN7SK) well-characterized RBP members of those respective ribonucleoprotein complexes. (d) Graph indicates accuracy (defined as (TP + TN) / (TP + TN + FP + FN)) for the top element (by relative information) from eCLIP for an RBP matching an annotated function for that RBP. Analysis was performed on 140 eCLIP datasets with functional annotation matching those listed in (c).

**Supplemental Fig. 3.**
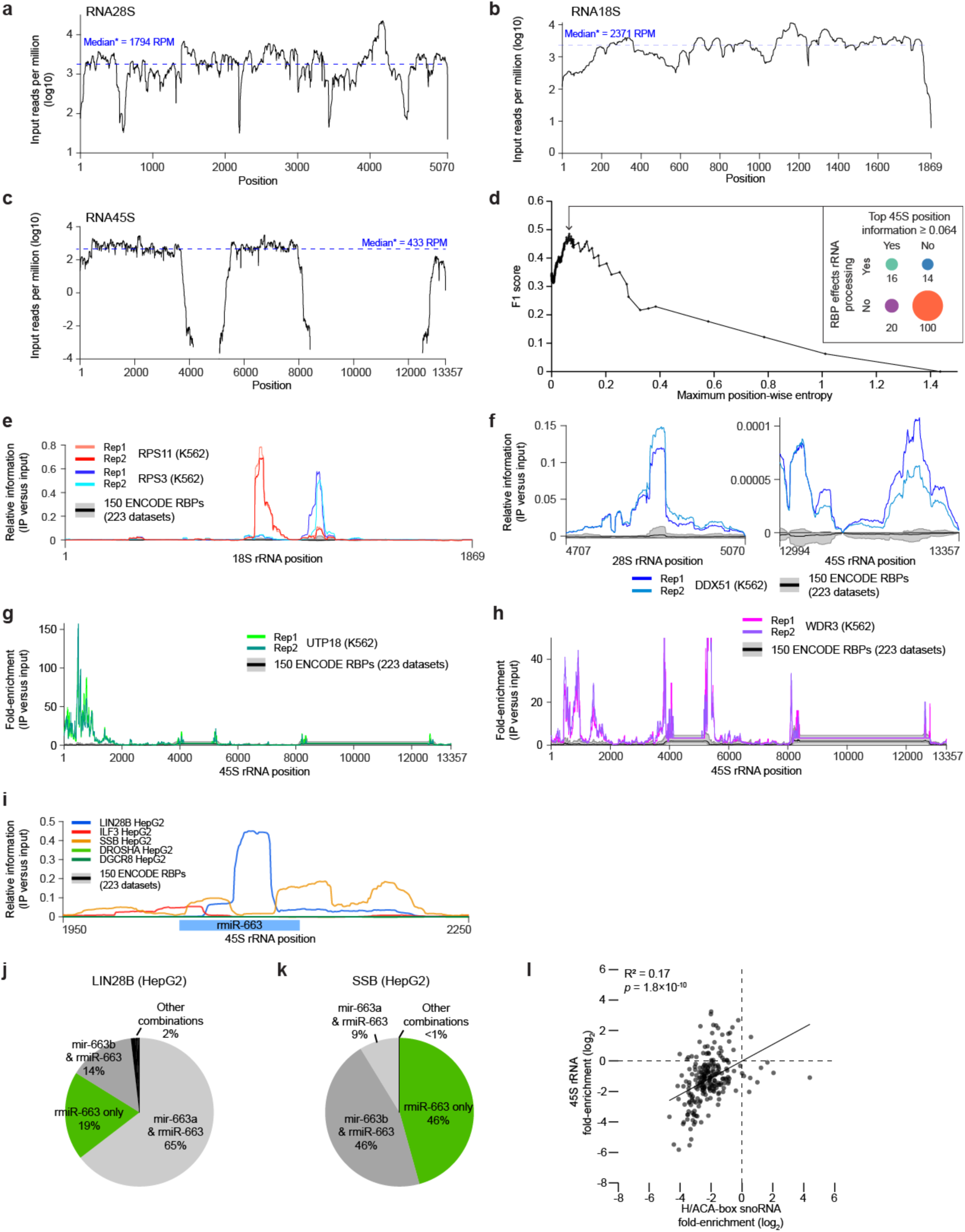
eCLIP enrichment for rRNA links RBPs with ribosomal RNA processing. (a-c) Line indicates average reads (per million non-PCR duplicate reads) mapping to each position across (a) 28S rRNA, (b) 18S rRNA, and (c) 45S rRNA. Blue line indicates median (calculated across all non-zero positions). Due to inability to distinguish between reads mapping to the 18S (or 28S) mature rRNA versus identical regions in the 45S precursor, reads mapping equally well to both are assigned to the 18S (or 28S), leading to gaps in the 45S plot. (d) Plot indicates (y-axis) F1 score for (x-axis) indicated relative information cutoffs, using RBPs with rRNA processing defects observed upon RNAi knockdown in Tafforeau *et al*. [27] as the ‘true positive’ reference set. (e-i) Lines indicate (y-axis) either fold-enrichment or relative information as indicated for (e) RPS11 in K562 and RPS3 in K562 for 18S rRNA, (f) DDX51 in K562 for 3’ end regions within (left) 28S rRNA and (right) 45S rRNA, (g) UTP18 in K562 for the 45S rRNA, (h) WDR3 in K562 for the 45S rRNA, and (i) indicated RBPs for the indicated 45S rRNA region flanking a putative ribosomal RNA-encoded microRNA (rmiR-663). For each, black line indicates mean and grey region indicates 10^th^ to 90^th^ percentile across all 223 eCLIP datasets. (j-k) Pie chart indicates for (j) LIN28B eCLIP in HepG2 or (k) SSB in HepG2 the fraction of reads that either map (green) with fewer mismatches to the rmiR-663 or (grey) to various combinations of rmiR-663 and genomic-encoded miR-663a/b and related family members (as shown in Fig. 3f). (l) Points indicate fold-enrichment in each eCLIP dataset for (x-axis) H/ACA-box snoRNAs versus (y-axis) the 45S rRNA precursor. Pearson correlation and significance was calculated in MATLAB.

**Sup. Fig. 4.**
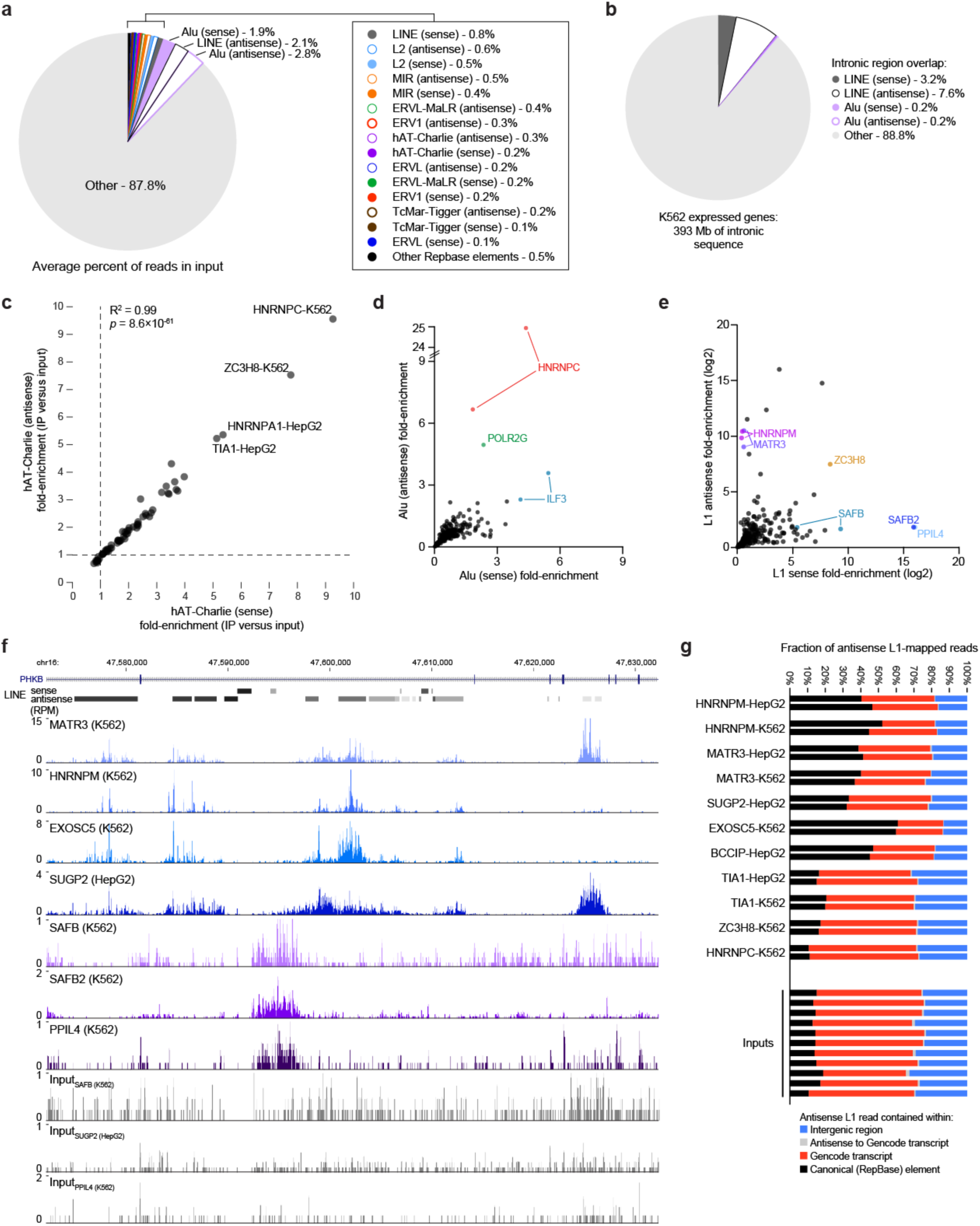
RBP association at retrotransposable and other repetitive elements. (a) Pie chart indicates the percent of reads (averaged across 223 eCLIP inputs) that map either uniquely to the genome within a RepeatMasker-predicted element, or are assigned to the consensus element sequencing in the family-aware mapping approach. Shown are RepBase element classes with average read density of at least 0.1%. Filled circles indicate sense elements, with open circles indicating antisense elements. (b) Pie chart indicates the fraction of intronic bases that overlap RepeatMasker-predicted L1 and Alu (sense and antisense) elements. Shown are genes with TPM≥1 in K562 cells, using the transcript isoform with the highest abundance in K562 rRNA-depleted RNA-seq. (c-e) Points indicate fold-enrichment (averaged across two biological replicates) for 223 eCLIP datasets for (c) hAT-Charlie, (d) Alu, and (e) L1 elements. In each, x-axis indicates sense and y-axis indicates antisense elements. Pearson R^2^ and significance was determined in MATLAB. (f) Genome browser image indicates read density for L1 sense- and antisense-enriched RBPs in intronic regions in *PKHB.* (g) Stacked bars indicate the fraction of antisense L1-assigned reads that were derived from mapping to (black) the canonical RepBase L1 element, (red) antisense L1 elements within Gencode v19 transcripts, (grey) to antisense L1 elements that are on the opposite strand from a Gencode v19 transcript, or (blue) antisense L1 elements in intergenic regions.

**Sup. Fig. 5.**
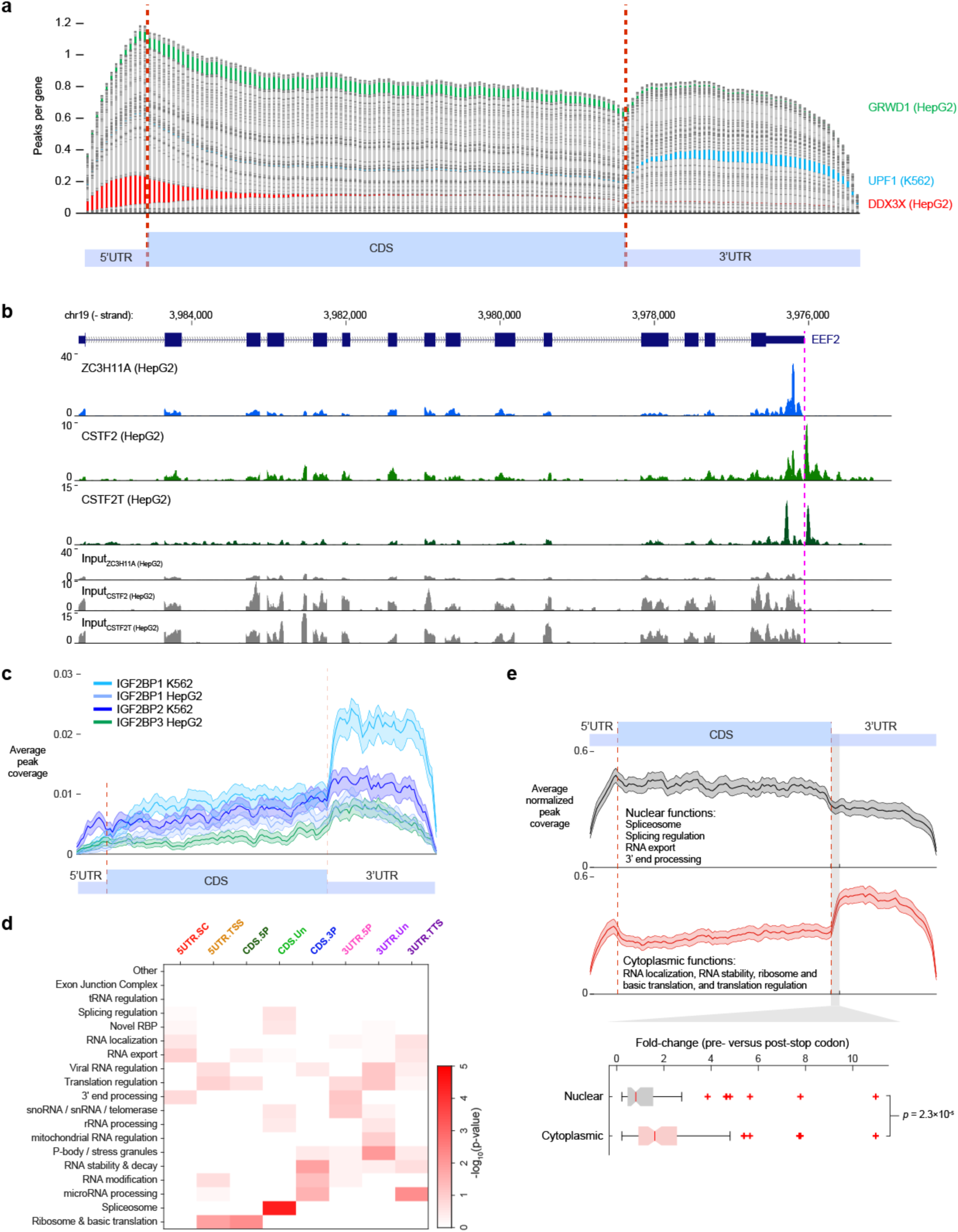
mRNA metagene profiles from eCLIP correspond to RBP RNA processing roles. (a) To create meta-mRNAs, each gene was normalized to 13 5’UTR, 100 CDS, and 49 3’UTR bins (based on average lengths among expressed transcripts in K562 cells). Stacked bars indicate the average number of significant and reproducible peaks for each RBP in each bin. (b) Tracks show read density in ZC3H11A, CSTF2, and CSTF2T eCLIP for *EEF2*. CSTF2 and CSTF2T show continuing read density past the annotated polyadenylation site. (c) Lines indicate average number of significant reproducible peaks per gene for bins across a meta-mRNA for IGF2BP family eCLIP. Shaded region indicates 5^th^ and 95^th^ percentile from 100 bootstrap samples. (d) Heatmap indicates significance (by Fisher’s Exact test, or Yates’ Chi-Square test where appropriate) of overlap between eCLIP datasets in indicated meta-mRNA cluster (x-axis) versus annotated RBP functions (y-axis). (e) (top) Lines indicate average normalized peak coverage for RBPs annotated with nuclear (*n* = 76) and cytoplasmic (*n* = 89) functions as indicated, with some RBPs present in both lists. Shaded region indicates standard error of the mean. (bottom) Box indicates 25^th^ to 75^th^ percentile (with median in red) of ratio between normalized peak coverage at bin 117 (post-stop codon) versus 113 (pre-stop codon). Significance was determined by two-sided Kolmogorov-Smirnov test.

**Sup. Fig. 6.**
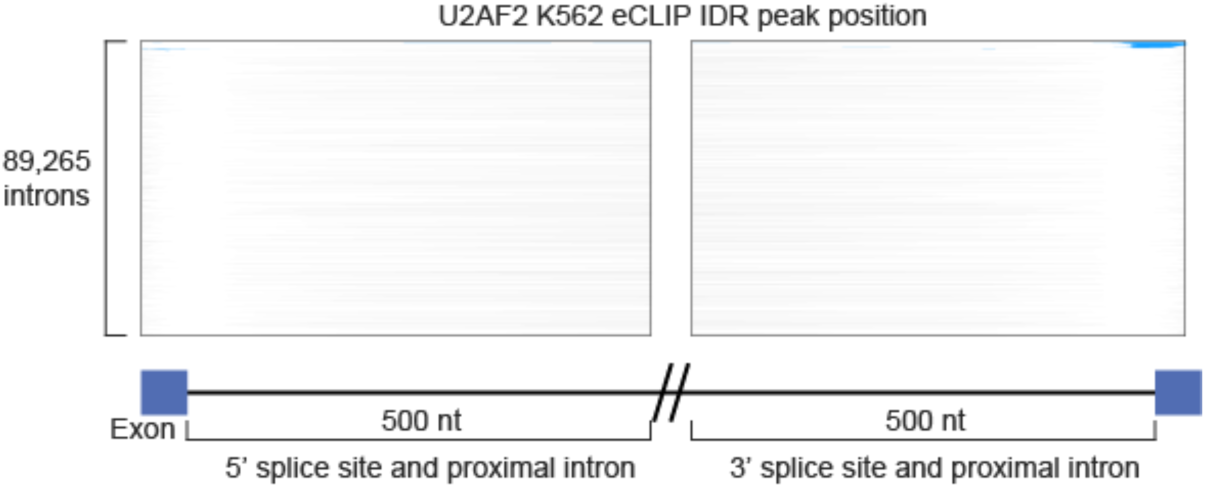
Meta-exon plots reveal intronic regulatory roles. Each line indicates the presence (in blue) of a reproducible U2AF2 K562 eCLIP peak for all 89,265 introns identified within genes with TPM ≥ 1 in K562 total RNA-seq. Region shown includes 500 nt of proximal intron and 50nt of exon flanking the 5’ and 3’ splice sites.

**Sup. Fig. 7.**
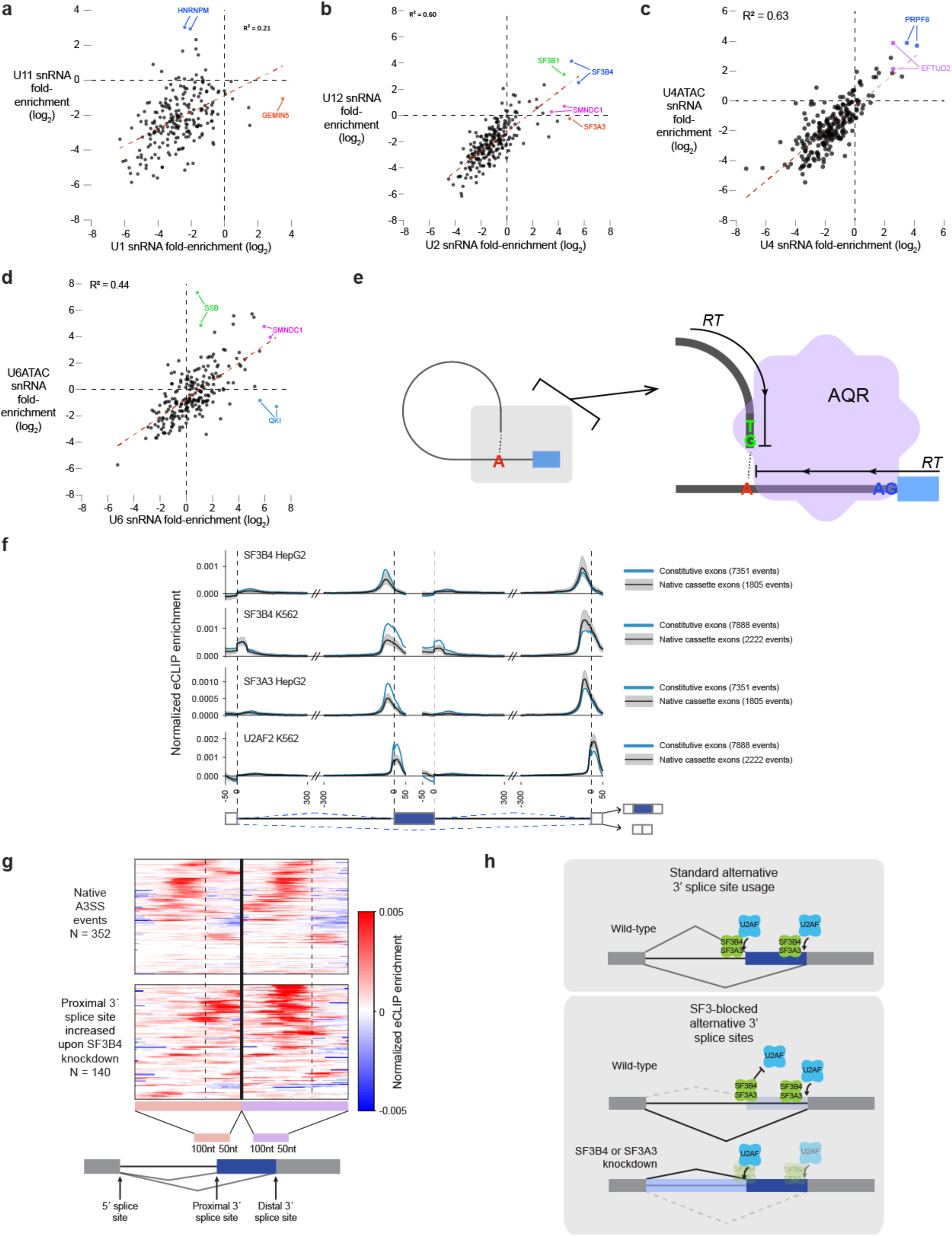
Insights from eCLIP of splicesome-associated RBPs. (a-d) Points indicate fold-enrichment in 223 eCLIP datasets for indicated (x-axis) major U2 spliceosomal snRNAs versus (y-axis) orthologous snRNAs in the U11/U12 minor spliceosome. (e) Model for AQR association with intronic lariats. Reverse transcription terminates at the lariat, generating reads with 5’ ends at the 5’ splice site as well as the position following the branch point adenosine. (f) Normalized splicing maps of SF3B4, SF3A3, and U2AF2 for (blue) constitutive exons versus (black) a set of ‘native’ cassette exons (nSE) with 0.05 < inclusion rate < 0.95 in controls. Lines indicate average eCLIP read density in IP versus input for indicated exon categories. Shaded area indicates 0.5th and 99.5th percentiles observed from 1000 random samplings of native events. The displayed region shown extends 50 nt into exons and 300 nt into introns. (g) Heatmap indicates normalized eCLIP signal for SF3B4 in HepG2 cells at alternative 3’ splice site events either (top) alternatively spliced in wild-type cells or (bottom) events with increased usage of the extended 3’ splice site upon SF3B4 knockdown. The region shown extends 50 nt into exons and 100 nt into introns. (h) Model for SF3B4 and SF3A3 blockage of 3’ splice site recognition by U2AF. At SF3-blocked alternative 3’ splice site events, knockdown of SF3 components leads to either usage of the upstream (proximal) 3’ splice site, or retention of the intron.

**Sup. Fig. 8.**
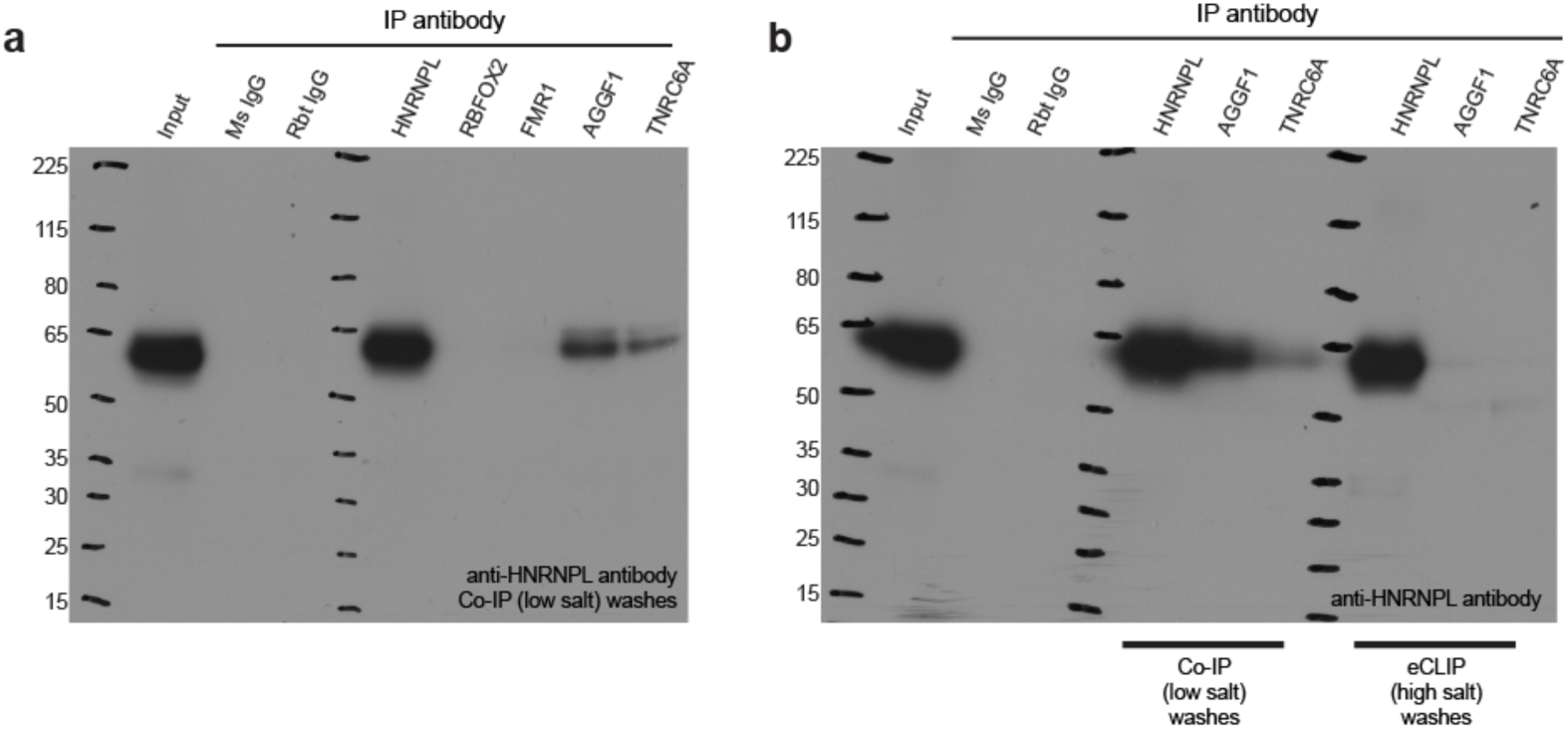
RBP co-association predicts known and novel RNP complexes. (a) Western blot for HNRNPL performed using 5 RBPs for immunoprecipitation (HNRNPL, RBFOX2, FMR1, AGGF1, and TNRC6A). Immunoprecipitation was performed using low-salt washes only. Also shown are immunoprecipitation using IgG isotype control for mouse and rabbit. (b) Western blot for HNRNPL using 3 RBPs for immunoprecipitation (HNRNPL, AGGF1, and TNRC6A), performed in two conditions: high salt washes (using standard eCLIP wash buffers), and low salt washes.

## Supplementary Tables

**Supplementary Table 1.** Accession identifiers for eCLIP datasets used in the manuscript.

**Supplementary Table 2.** RNA binding protein function annotations, localization patterns, and predicted RNA binding domains.

**Supplementary Table 3.** List of multi-copy element annotations used in family-aware mapping.

**Supplementary Table 4.** Quantitation of multi-copy RNA family enrichment for 223 eCLIP datasets.

## Declarations

### Ethics approval and consent to participate

Not applicable

### Consent for publication

Not applicable

### Availability of data and materials

Raw and processed eCLIP data is available at the ENCODE Data Coordination Center (https://www.encodeproject.org). Accession identifiers for datasets used are provided in Supplementary Table 1.

### Competing interests

ELVN is co-founder, member of the Board of Directors, on the SAB, equity holder, and paid consultant for Eclipse BioInnovations. GWY is co-founder, member of the Board of Directors, on the SAB, equity holder, and paid consultant for Locana and Eclipse BioInnovations. The terms of these arrangements have been reviewed and approved by the University of California, San Diego in accordance with its conflict of interest policies. The authors declare no other competing financial interests.

### Funding

This work was funded by the National Human Genome Research Institute ENCODE Project, contract U54HG007005 to BRG (principal investigator) and GWY (co-principal investigator), and U41HG009889 to BRG (PI) and GWY (PI). ELVN is a Merck Fellow of the Damon Runyon Cancer Research Foundation (DRG-2172-13) and is supported by a K99 grant from the NHGRI (HG009530).

### Authors’ contributions

ELVN, SMB, JM, SP, KEG, CGB, TBN, IR, RS, BS, and RW generated eCLIP and RNP visualization data. ELVN, GAP, and BAY performed data analysis and software development. ELVN, XDF, BRG, and GWY wrote the paper and led data generation and analysis.

## Acknowledgements

We thank members of the Yeo lab, as well as Christopher Burge, Eric Lécuyer, and Stefan Aigner, and members of the Graveley, Burge, Lécuyer, and Fu labs for helpful comments and suggestions during the development of this work.

## Notes

#### Summary of Updates

added supplemental tables

